# Remodeling of Crossbridges Controls Peptidoglycan Cross-linking Levels in Bacterial Cell Walls

**DOI:** 10.1101/840405

**Authors:** Sean E. Pidgeon, Alexis J. Apostolos, Marcos M. Pires

**Author notes:** Authors contributed equally to this work (Sean E. Pidgeon, Alexis J. Apostolos).

## Abstract

Cell walls are barriers found in almost all known bacterial cells. These structures establish a controlled interface between the external environment and vital cellular components. A primary component of cell wall is a highly crosslinked matrix called peptidoglycan (PG). PG crosslinking, carried out by transglycosylases and transpeptidases, is necessary for proper cell wall assembly. Transpeptidases, targets of β-lactam antibiotics, stitch together two neighboring PG stem peptides (acyl-donor and acyl-acceptor strands). We recently described a novel class of cellular PG probes that were processed exclusively as acyl-donor strands. Herein, we have accessed the other half of the transpeptidase reaction by developing probes that are processed exclusively as acyl-acceptor strands. The critical nature of the crossbridge on the PG peptide was demonstrated in live bacterial cells and surprising promiscuity in crossbridge primary sequence was found in various bacterial species. Additionally, acyl-acceptor probes provided insight into how chemical remodeling of the PG crossbridge (e.g., amidation) can modulate crosslinking levels, thus establishing a physiological role of PG structural variations. Together, the acyl-donor and -acceptor probes will provide a versatile platform to interrogate PG crosslinking in physiologically relevant settings.

**SYNOPSIS TOC:** 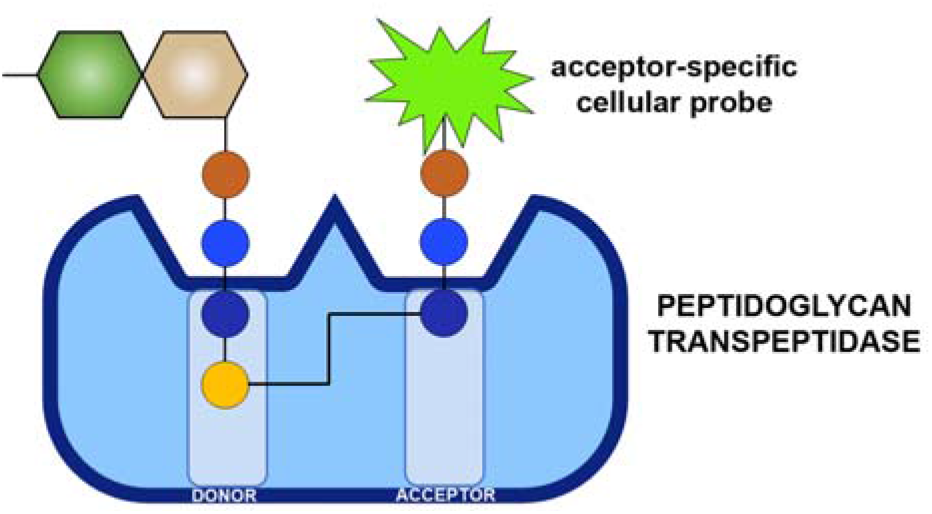

---

Gram-positive pathogens continue to impose a significant global public health burden.^1^ Among these organisms, Enterococci are one of the leading causes of nosocomial infections.^2^ With the rise in antibiotic drug resistance, it has become clear that more bacterial targets are needed to improve the therapeutic options to fight Enterococci infections. Peptidoglycan (PG), an essential component of bacterial cell walls, is one of the most important targets of antibiotics. Disaccharides of *N*-acetylglucosamine (GlcNAc) and *N*-acetylmuramic acid (MurNAc) make up the basic PG unit, which is further decorated with a pentameric stem peptide (e.g., L-Ala-D-isoGlx-L-Lys-D-Ala-D-Ala).^3–4^ PG units are polymerized and crosslinked *via* transglycosylation and transpeptidation (TP) reactions that assemble the sugars and peptides, respectively. PG crosslinking is essential, and, therefore, molecules that block this reaction are traditionally valuable antibiotics (e.g., β-lactams and vancomycin) and continue to result in promising drug leads (e.g., teixobactin).^5–7^

Intense efforts have led to a greater understanding of the chemical makeup, mechanism of assembly, and essential proteins that construct the PG network. Moreover, there is some evidence that small structural modifications within the PG matrix can alter the overall plasticity and physical properties of the cell wall (e.g., resistance to lysozyme).^8–10^ Chemical composition of the stem peptide primary sequence can vary by the amidation/methylation of carboxylic groups, removal of amino acids, variation of amino acid character, and extensive alteration to the central lysine group. Yet, for the most part, their physiological impact remains elusive.^11–14^

The 3^rd^ position with the PG stem peptide has defining structural characteristics. As an example, the difference of a carboxylic acid group on L-Lys verus *meso*-2,6-diaminopimelic acid (*m*-DAP) has important implications in the activation of the human innate immune system.^15^ In *Enterococcus faecium* (*E. faecium*), the lysine sidechain is modified with D-iAsx while *Enterococcus faecalis* (*E. faecalis*) displays L-Ala-L-Ala. In addition, the nature of the crossbridge (the amino acids anchored onto the lysine residues) is critical based on its role in the nucleophilic step during crosslinking reactions. There are two main classes of PG cross-linking enzymes: D,D-TPs and L,D-TPs (Figure 1). Crosslinking by Penicillin Binding Protein (PBP) D,D-TPs generate 4-3 peptide crosslinks and L,D-TPs (Ldts) create 3-3 crosslinks within PG.

**Figure 1.**
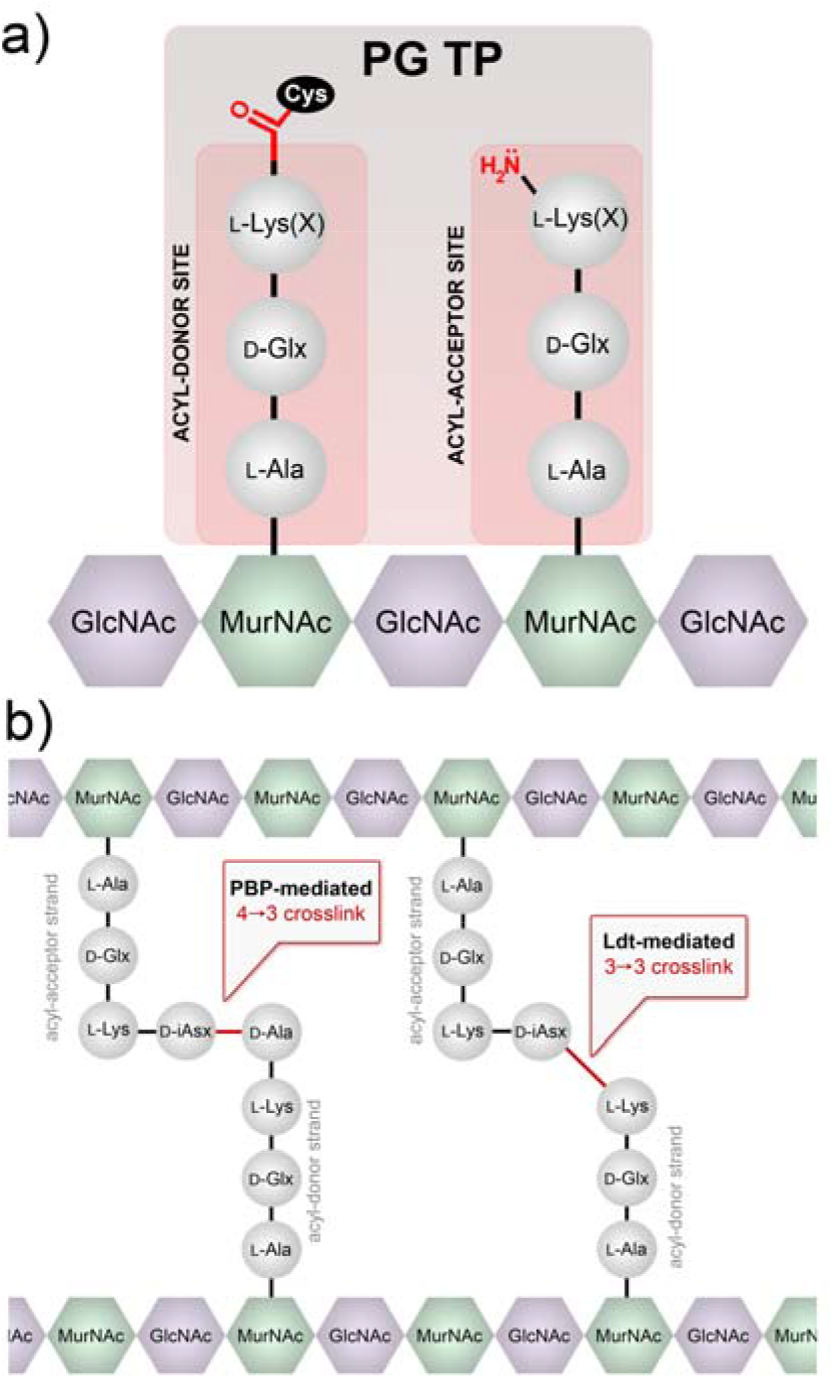
Crosslinking reaction by TPs. (a) Example of two binding sites (acyl-donor and -acceptor) at the intermediate step of the crosslinking reaction are depicted. The acyl-chain is covalently anchored on the TP, which is subsequently transferred to the amino group on the acyl-acceptor strand. X represents the crossbridging amino acids. (b) Schematic of PG crosslinking in *E. faecium* showing the 2 types of crosslinks observed.

Fundamental understanding of PG crosslinking is key to define the processes that control PG structure and may provide novel drug targets. Single D-amino acid PG probes have spurred valuable insight into PG crosslinking in live bacterial cells.^16–21^ Furthermore, elegant prior *in vitro* studies using synthetic PG analogs have provided insight into the substrate structural requirements of TPs.^12, 22–25^ However, fewer reports have described probes that decipher TP substrate requirements in live bacterial cells. A recent example of a synthetic L-Lys-D-Ala-D-Ala stem peptide mimic led to the demonstration of extraseptal TP activity in *Staphylococcus aureus* (*S. aureus*).^26^ Likewise, we recently reported the use of synthetic PG probes to establish crosslinking parameters based on the acyl-donor strand of Ldts.^27^ We have now built on those findings by developing complementary probes that selectively track and dissect acyl-acceptor strand processing by TPs in live bacterial cells.

We initially hypothesized that structural analogs of the two TP substrates could be developed to control active site loading. In designing our previous acyl-donor probe **1**, we mimicked the stem peptide fragment as a tetrapeptide and blocked the nucleophilic site on the acyl-acceptor strand (Figure 2a). In this work, the acyl-acceptor specific probe **2** was designed by installing the acyl-acceptor fragment (crossbridging D-iAsx in the case of *E. faecium*) and removing the terminal fragment recognized by the acyl-donor site (terminal D-Ala and/or D-Ala-D-Ala). Therefore, PG probes with the basic scaffold of tripeptide **2** should act solely as an acyl-acceptor strand. To track incorporation into the PG, a fluorescent handle was added to the *N*-terminus of the stem tripeptide analogs. Incubation of live bacterial cells with tripeptide probes is expected to result in their covalent incorporation into the PG scaffold by TPs (Figure 2b). Crosslinking can then be readily quantified and analyzed using standard fluorescence-based techniques, thus providing a mode to interrogate how primary sequences of the acyl-acceptor strand modulate PG crosslinking (Figure 2c).

**Figure 2.**
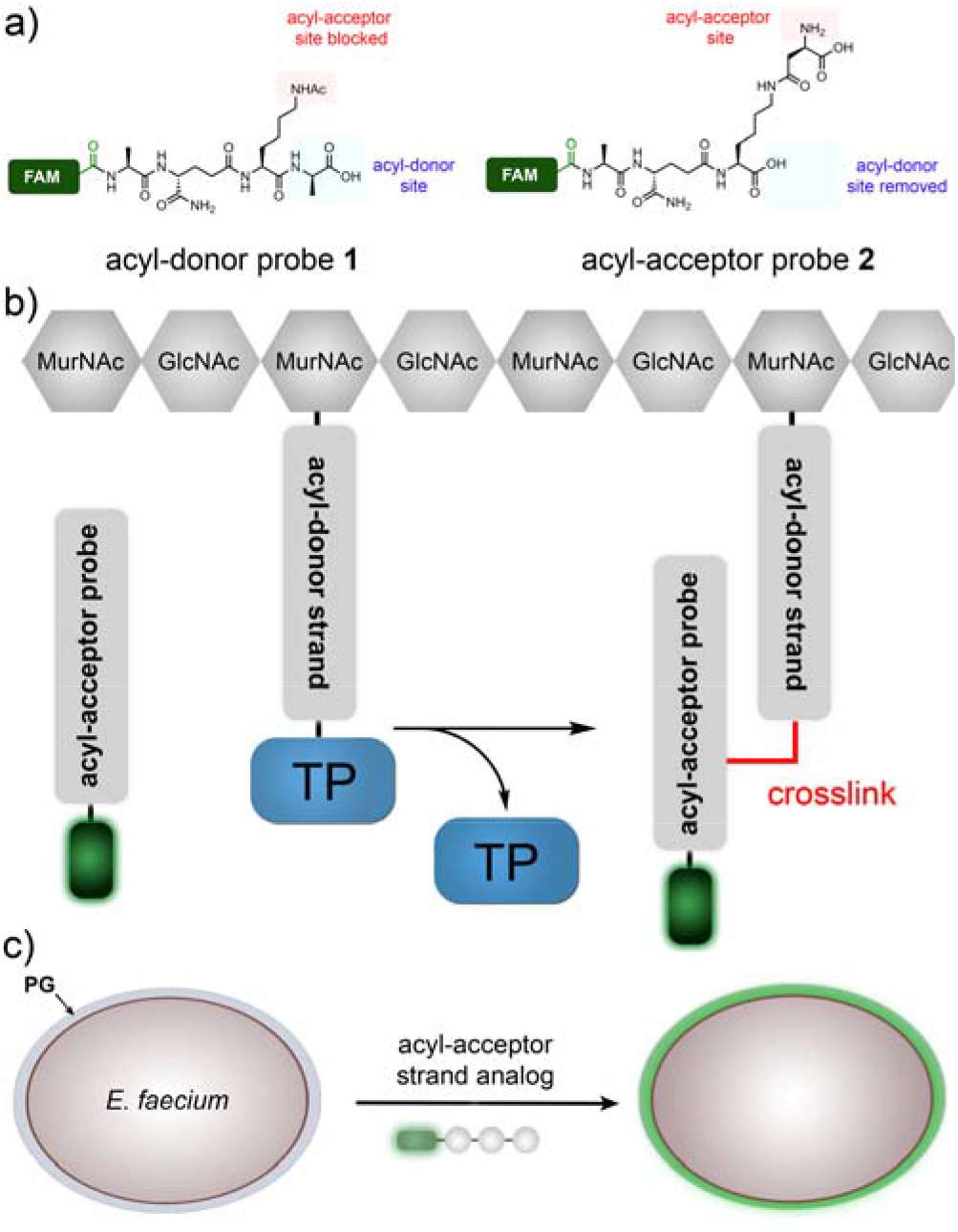
Design of tripeptide probes and the assay to assess acyl-acceptor strand in live cells. (a) Structure-based design of acyl-donor and -acceptor probes. (b) Schematic diagram showing how the fluorescently tagged tripeptide probe gets covalently crosslinked into the bacterial PG scaffold. (c) Schematic of the whole cell analysis that can be accessed by the tripeptide probes.

At first, we synthesized a panel of tripeptide probes that were designed to investigate acyl-acceptor strand recognition in live *E. faecium* (Figure 3). D-iAsx is the canonical crossbridge amino acid in *E. faecium*, where a nucleophilic attack by the sidechain *N*-terminus nitrogen leads to formation of PG crosslinks. All probes were synthesized using standard solid phase peptide chemistry and included a carboxy-fluorescein tag (FAM). *E. faecium* cells were treated with individual probes and cellular fluorescence levels were measured after overnight incubation. The first two stem peptide mimics, **TriQK** and **TriQD**, were designed to test the role of crossbridging amino acids in the acyl-capture step of TP. Our results showed that when cells were treated with **TriQK**, which lacks the crossbridging amino acid, fluorescence levels were near background (Figure 3a). Introduction of the crossbridge D-iAsp (**TriQD**) led to ~7-fold fluorescence increase over background. These results provide *in vivo* evidence for the importance of the crossbridge for robust crosslinking of the PG scaffold and suggest that the enzyme responsible for D-iAsp installation, Asl_fm_,^28^ is a potential narrow-spectrum drug target. Given that only approximately 60% of the lysines are modified with D-iAsx in *E. faecium*^29^, and that we have evidence that unmodified lysine residues do not participate in TP reactions as acyl acceptors, these results also suggest a mode of controlling PG crosslinking based on Asl_fm_ processing of PG precursors in the cytosol of cells prior to exposure to TPs.

**Figure 3.**
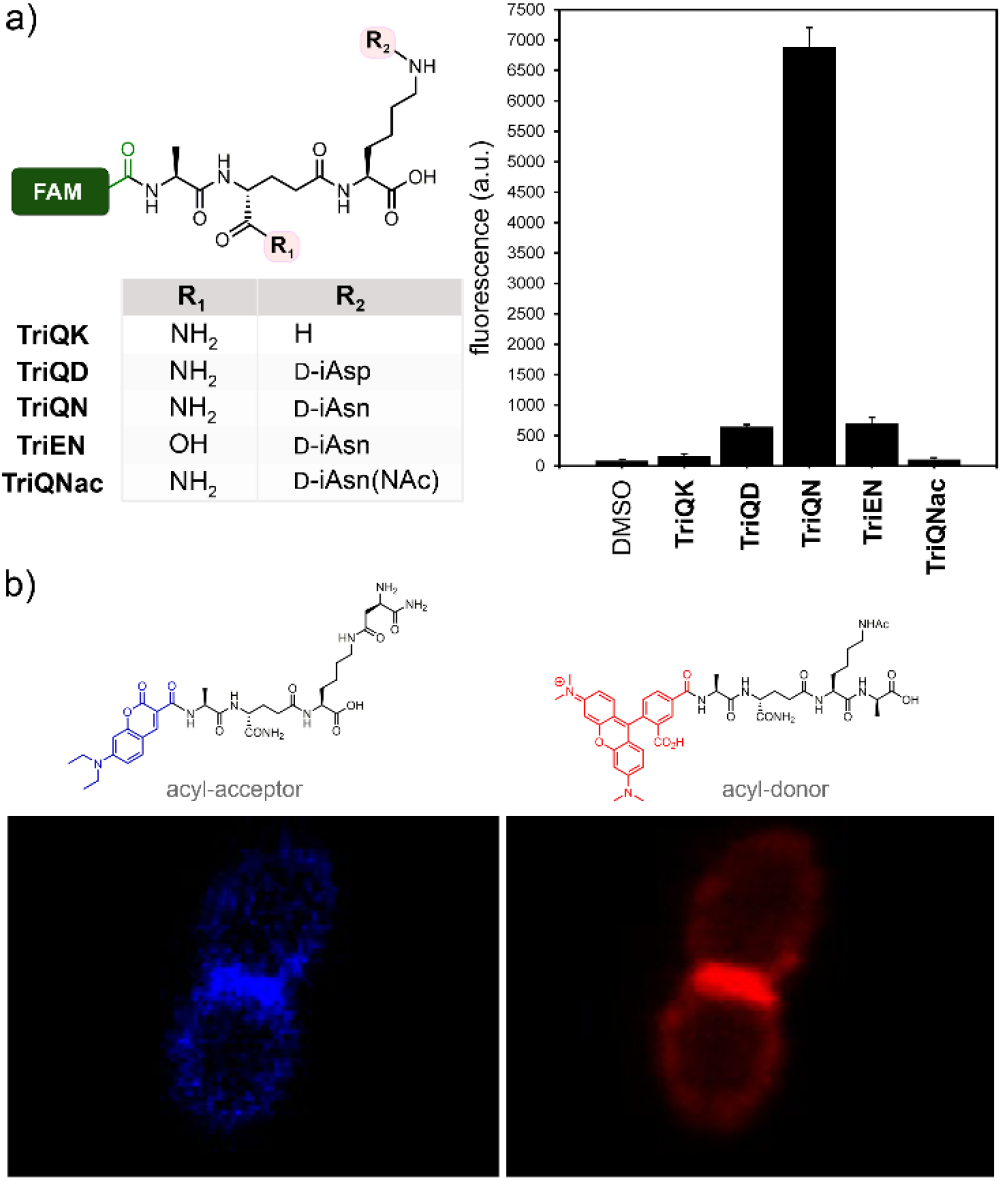
Crosslinking of tripeptide probes in live *E. faecium* cells. (a) Chemical series tripeptide probes based on the modifications at the D-iGlx and lysine sidechain; FAM = carboxyfluorescein. Flow cytometry analysis of *E. faecium* (D344s) treated overnight with 100 µM of tri-peptide probes. Data are represented as mean + SD (n = 3). (b) Confocal microscopy of *E. faecium* cells labeled with the designated probes (100 μM) for 5 min. Scale bar = 2 μm.

We next evaluated how chemical remodeling of the acyl-acceptor strand may influence TP crosslinking in *E. faecium*. While the sidechain of the 3^rd^ position lysine is initially loaded with a D-iAsp crossbridge, its carboxylic acid can be biochemically amidated to generate D-iAsn crossbridges to varying levels in *E. faecium*.^29^ The physiological and TP-processing consequences of D-iAsp amidation remain unresolved due to confounding chemical re-arrangements (e.g., deamidation of D-iAsn and cyclization) during PG isolation and analysis.^29^ Instead, we hypothesized that we could establish the role of amidation using synthetic analogs in live bacteria. Strikingly, treatment of *E. faecium* with **TriQN** resulted in considerably higher crosslinking levels relative to **TriQD** (Figure 3). These results suggest that amidation of D-iAsp within the acyl-acceptor strand play a critical role in controlling PG crosslinking levels.

We then analyzed the role of amidation in the 2^nd^ position D-iGlu in the crosslinking step. Recent genetic analyses revealed the essential nature of the genes encoding the enzymes that carry out the amidation of D-iGlu to D-iGln in *S. aureus* and *Streptococcus pneumoniae*.^30–31^ Subsequent *in vitro* characterization suggested that lack of D-iGlu amidation impairs PG crosslinking.^11, 23^ Yet, it remained unresolved whether D-iGlu amidation impacts processing at the acyl-donor, acyl-acceptor, or both sites. We recently demonstrated that amidation of D-iGlu is critical for the acyl-donor strand for both classes of TPs.^27^ To test these concepts in the acyl-acceptor substrate, *E. faecium* cells were treated with **TriEN**. In sharp contrast to the D-iGln variant (**TriQN**), cellular fluorescence levels were significantly lower in cells treated with the D-iGlu variant (**TriEN**). Together, our data point to a three-level biochemical control (addition of crossbridge, D-iGlu to D-iGln, and D-iAsp to D-iAsn) of the acyl-acceptor substrate structure that likely combine to tune cross-linking level in *E. faecium*.

To confirm the incorporation of **TriQN** into the PG scaffold, metabolically labeled PG were digested and the resulting fragments analyzed (data not shown). Moreover, we demonstrated that the crossbridge of **TriQN** is, in fact, the acyl-acceptor site by building a control probe (**TriQNAc**) in which the nucleophilic amino group is acetylated to block acyl-capture. Fluorescence levels for cells treated with **TriQNAc** were near background, a result that is consistent with the site of acyl-capture. Confocal microscopy was then used to delineate the incorporation of the probes within the PG scaffold of live cells. For these experiments, a short pulse labeling step was performed with the acyl-acceptor probe modified with a blue fluorophore and the acyl-donor probe modified with a red fluorophore (Figure 3b). From our results, it is clear that there is some overlap between PG incorporation of these probes and we established that the septal region is the primary site of PG incorporation.

To extend our results to another type of Enterococci, acyl-acceptors probes were adopted for *E. faecalis*. In the case of *E. faecalis*, ligases BppA1 and BppA2 are responsible for transferring L-Ala from Ala-*t*RNA to the first and 2^nd^ positions of the ε-amino group of lysine of PG precursors, respectively.^32^ We synthesized **TriQAA**, which mimics the crossbridge in *E. faecalis*, to initially test acyl-acceptor processing (Figure 4a). Incubation of *E. faecalis* cells with **TriQAA** led to a ~10.5 fold increase over the unmodified lysine side chain **TriQK**, thus again demonstrating the necessity for a crossbridging unit for proper acyl capture strand in Enterococci PG crosslinking. Inverting the stereochemistry of the crossbridge in **TriQAA** to **TriQaa** also led to baseline fluorescence, indicating that TPs are stereospecific at the acyl-acceptor position. Acetylation of the *E. faecalis* probe, **TriQAAac**, which blocks acyl-acceptor nucleophilic site, led to baseline fluorescence levels. This finding is consistent with the amino group of the terminal L-alanine as the nucleophile for covalent crosslinking into the PG scaffold. Surprisingly, lack of amidation at the second position (**TriEAA**) only resulted in a ~1.3-fold decrease in fluorescence, showing that amidation may not be an absolute requirement for the acyl-acceptor recognition in *E. faecalis*.

**Figure 4.**
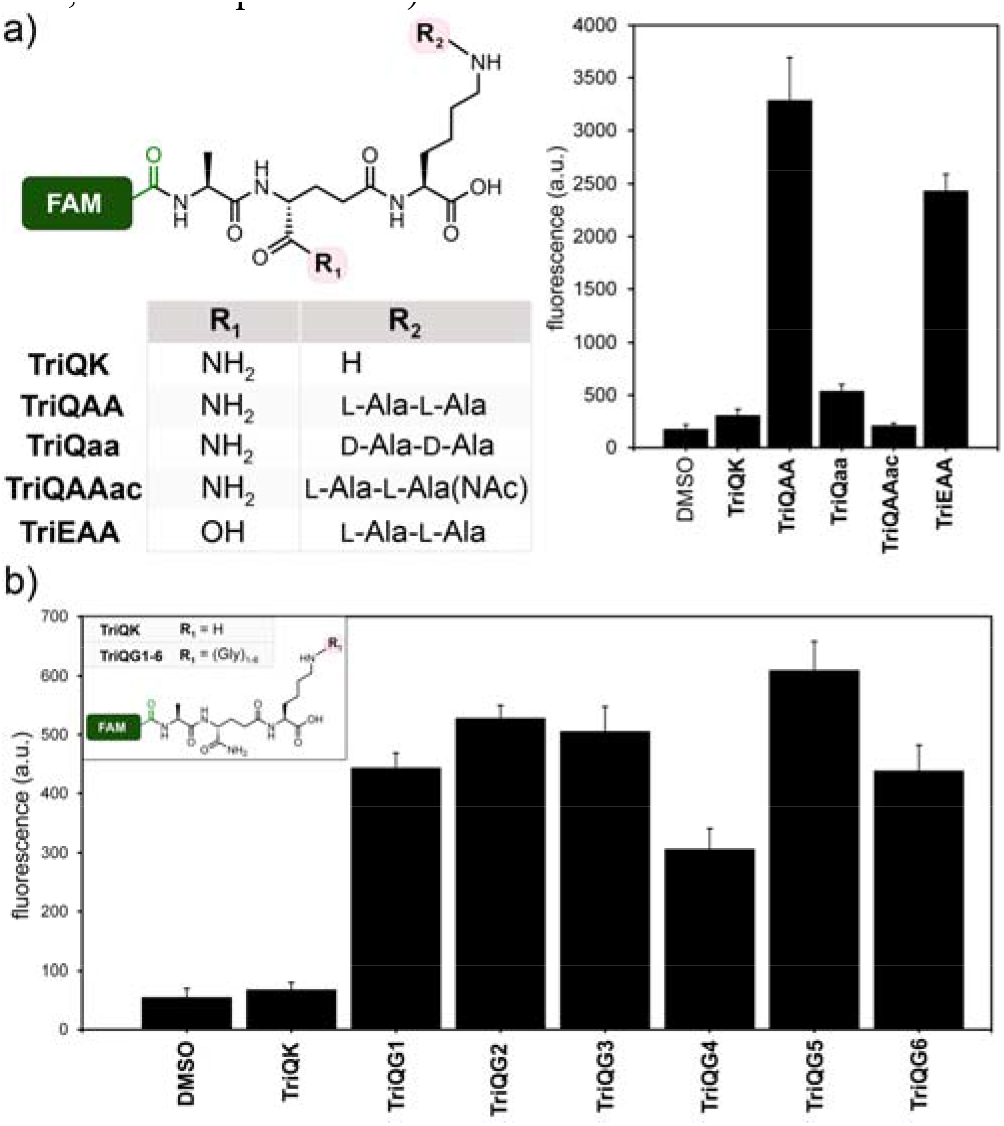
Crosslinking of tripeptide probes in live *E. faecalis* and *S. aureus* cells. Chemical series tripeptide probes based corresponding to the stem peptide of (a) *E. faecalis* and (b) *S. aureus*. Flow cytometry analysis of cells treated overnight with 100 µM of tri-peptide probes. Data are represented as mean + SD (n = 3).

We next turned our attention to the problematic human pathogen *S. aureus*. In *S. aureus*, peptidyl transferases FemX, FemA, and FemB catalyze the stepwise addition of five glycines to the lysine residue on the 3^rd^ position of the PG precursor.^33–35^ Recently, Walker and co-workers isolated PG precursors corresponding to one, three, and five glycine units and showed *in vitro* that PBP transpeptidases have varying levels of preference for the length of the glycine chain.^12^ We built a library of tripeptide *S. aureus* probes that complement their approach by assessing the acyl-acceptor preference in live bacterial cells. We synthesized **TriQG1-6** (Figure 4b), corresponding to glycine chains from 1 to 6 units, and measured their incorporation into the PG scaffold of *S. aureus* cells. Consistent with our previous results, the absence of glycine modification (**TriQK**) resulted in background levels of PG incorporation. Interestingly, addition of a single glycine (**TriQG1**) led to ~7-fold increase in cellular fluorescence levels. These results are consistent with a previously reported viable *S. aureus femAB*-null strain in which one glycine crossbridges were observed, albeit with severe crosslinking defects and impaired growth.^35–36^ Our results clearly demonstrate the physiological consequence of a lack of a single glycine unit and point to FemX is an particularly attractive target for therapeutic development, something that had been suggested but had not been observed in live bacterial cells due to the lethality of *femX*-deletion.^37^ Crossbridges of 2 to 5 glycine resulted in comparable labeling levels. Extending beyond the natural pentaglycine chain (**TriQG6**) did not significantly impact crosslinking, which suggests that there is a high level of flexibility by the acyl-acceptor strand in *S. aureus*.

Having individually established the structural preference of the acyl-acceptor strand for various bacteria, we next set out to examine how crossbridges are tolerated by TPs across different species. Because TPs are frequent targets of β-lactams and other potent antibiotics, these enzymes play central roles in the drug resistance among several classes of medically relevant bacteria.^38^ Moreover, mobile elements carrying TP genetic information can be a powerful modality for the acquisition of drug resistant phenotypes. The *mecA* gene underpinning the drug resistant phenotype of Methicillin-resistant *Staphylococcus aureus* (MRSA) has been proposed to originate from *Staphylococcus sciuri* (*S. sciuri*).^39^ Therefore, the ability of TPs to process diverse PG architectures may have implications on the potential of TP to cross polinate drug resistance phenotypes. As an example, it was recently shown that commensal streptococci can serve as a reservoir for PBPs that can cross pollinate drug resistant phenotypes.^40^

To investigate the tolerance of non-native crossbridges in PG crosslinking reactions, the three canonical crossbridges (**TriQN**, **TriQAA**, and **TriQG5**) were incubated with *E. faecium*, *E. faecalis*, and *S. aureus*. *S. aureus* showed some promiscuity in accepting both **TriQAA** and **TriQN** (Figure 5 and **Figure SX** for raw values) while still demonstrating some preference for pentaglycine crossbridge. Conversely, *E. faecium* showed strong preference for its natural acyl-acceptor strand **TriQN**. Treatment of *E. faecium* cells with **TriQAA** or **TriQG5** led to background fluorescence levels. Strikingly, *E. faecalis* showed little preference in the acyl-acceptor side chain and labeled well with all three tripeptide probes.. These findings are consistent with previous *in vitro* studies that illustrated the promiscuity of *E. faecalis* TPs in accepting non-native crossbridges.^41–42^ Our results provide *in vivo* evidence that some bacterial strains possess a high degree of tolerance for exogenous crossbridges, which may ultimately provide a potentially viable drug resistance modality.

**Figure 5.**
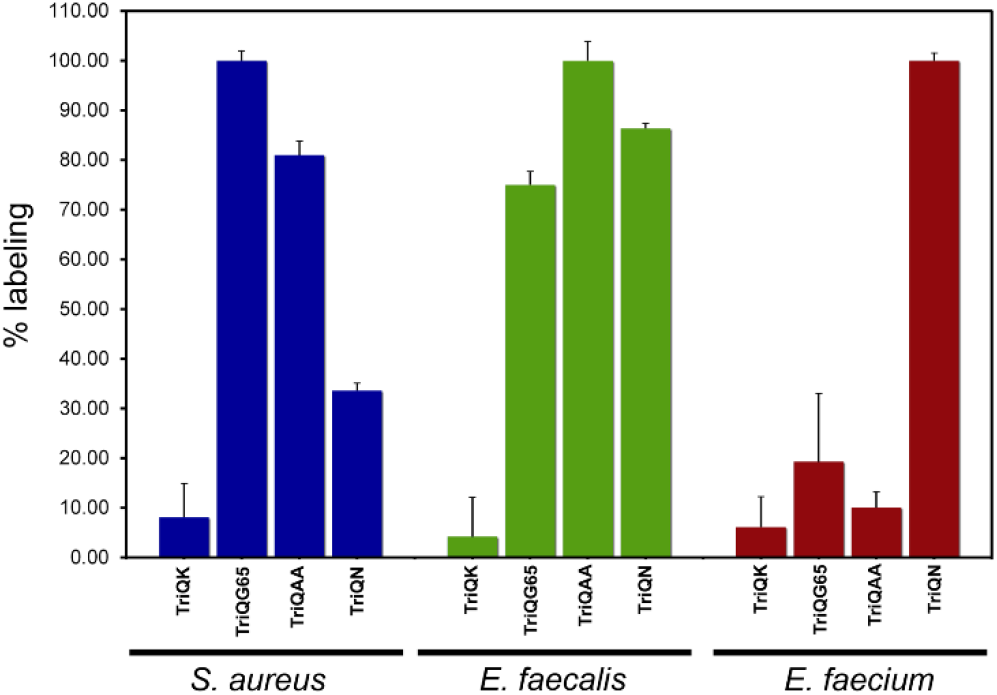
Crossbridge preferences across bacterial species. Flow cytometry analysis of cells treated overnight with 100 µM of tripeptide probes. Data are represented as mean + SD (n = 3).

Targeting the enzymatic processes that control PG assembly is a powerful method of disabling bacterial pathogens. Overall, our results demonstrate the requirement of the crossbridge in acyl-acceptors for proper crosslinking by TPs in three different types of bacteria, thus reinforcing the concept that lysine modification is pivotal in tuning the PG crosslinking machinery. Future investigations will center on finding out how flexibility in PG crosslinking may promote the evolution of drug resistance. In developing a comprehensive structural analysis of both the acyl-donor and - acceptor strands, we anticipate that these results may guide drug design and open new avenues of therapeutic modalities.

## ASSOCIATED CONTENT

### Supporting Information

Additional experimental details (methods, characterization and synthesis of peptide probes) and figures.

## AUTHOR INFORMATION

### Notes

The authors declare no competing financial interests.

## ACKNOWLEDGMENT

This study was supported by GM124893-01 (M.P).

