## supporting for "Remodeling of Crossbridges Controls Peptidoglycan Cross-linking Levels in Bacterial Cell Walls"

**Supporting Information**

**Materials**. All peptide related reagents (resin, coupling reagent, deprotection reagent, amino acids, and cleavage reagents) were purchased from ChemImpex. Bacterial strains *E. faecium* D344s and M9 and *E. faecalis* ATCC 29212 were grown in brain heart infusion broth for all experiments. *S. aureus* SC01 was grown in lysogeny broth.

**Flow cytometry analysis of bacteria labeling with *E. faecium*.** Brain heart infusion (BHI) broth containing 100 µM **Fl-Tri(Lys), Fl-Tri(Asn), Fl-Tri(Asp),** or **Fl-Tri-Glu(Asn)** were prepared. *E. faecium* D344s or *E. faecium* M9 from an overnight culture were added to the medium (1:100 dilution) and allowed to grow overnight at 37 ^o^C with shaking at 250 rpm. The bacteria were harvested at 6,000g and washed three times with original culture volume of 1x PBS followed by fixation with 2% formaldehyde in 1x PBS for 30 min at ambient temperature. The cells were washed once more to remove formaldehyde and then analyzed using a BDFacs Canto II flow cytometer using a 488nm argon laser (L1) and a 530/30 bandpass filter (FL1). A minimum of 10, 000 events were counted for each data set. The data was analyzed using the FACSDiva version 6.1.1.

**Flow cytometry analysis of bacteria labeling with *E. faecalis*.** Brain heart infusion (BHI) broth containing 100 µM **Fl-Tri(Lys), Fl-Tri(L-Ala-L-Ala), Fl-Tri(D-Ala-D-Ala),** or **Fl-Tri(L-Ala-L-Ala-Ac)** were prepared. *E. faecalis* ATCC 29212 from an overnight culture was added to the medium (1:100 dilution) and allowed to grow overnight at 37 ^o^C with shaking at 250 rpm. The bacteria were harvested at 6,000g and washed three times with original culture volume of 1x PBS followed by fixation with 2% formaldehyde in 1x PBS for 30 min at ambient temperature. The cells were washed once more to remove formaldehyde and then analyzed using a BDFacs Canto II flow cytometer using a 488nm argon laser (L1) and a 530/30 bandpass filter (FL1). A minimum of 10, 000 events were counted for each data set. The data was analyzed using the FACSDiva version 6.1.1.

**Flow cytometry analysis of bacteria labeling with *S. aureus*.** Lysogeny broth (LB) broth containing 100 µM **Fl-Tri(Lys), Fl-Tri(Gly_1_), Fl-Tri(Gly_2_), Fl-Tri(Gly_3_), Fl-Tri(Gly_4_), Fl-Tri(Gly_5_),** or **Fl-Tri(Gly_6_)** were prepared. *S. aureus* SC01 from an overnight culture was added to the medium (1:100 dilution) and allowed to grow overnight at 37 ^o^C with shaking at 250 rpm. The bacteria were harvested at 6,000g and washed three times with original culture volume of 1x PBS followed by fixation with 2% formaldehyde in 1x PBS for 30 min at ambient temperature. The cells were washed once more to remove formaldehyde and then analyzed using a BDFacs Canto II flow cytometer using a 488nm argon laser (L1) and a 530/30 bandpass filter (FL1). A minimum of 10, 000 events were counted for each data set. The data was analyzed using the FACSDiva version 6.1.1.

**Confocal microscopy analysis of *E. faecium* labeled with TetraRh, PentaFl, and DADA.** Brain heart infusion (BHI) broth containing 500 µM **TetraRh,** 500 µM **PentaFl,** and 5 mM **DADA** was prepared. *E. faecium* (D344s) from an overnight growth was added to the medium (1:10 dilution) and incubated at 37 ^o^C with shaking at 250 rpm for 5 minutes. The bacteria were immediately harvested at 6,000g and washed three times with original culture volume of 1x PBS followed by fixation with 2% formaldehyde in 1x PBS for 30 min at ambient temperature. The cells were washed once more to remove formaldehyde and then analyzed using a Nikon C2 confocal microscope.

Peptidoglycan Isolation. BHI medium (200 mL) containing 500 µM Fl-Tri(Asp) was prepared. *E. faecium* D344s bacteria were added to the BHI medium (1:100) and allowed to grow overnight at 37 ^o^C with shaking at 250 rpm. The cells were harvested and washed with 1x phosphate buffer saline (PBS) (3 × 50 mL each). The cells were then resuspended in 1x PBS and boiled for 7 min and then centrifuged at 14,000g for 8 min at 4 °C. Cells were then placed in 25 mL of 5% (w/v) sodium dodecyl sulfate (SDS) and boiled for 25 min followed by centrifugation at 14,000g for 8 min at 4 °C. Following centrifugation, cells were boiled again in 25 mL of 4% (w/v) SDS for 15 min followed by centrifugation using same parameters as before. Cells were then washed 5 times with 60 °C DI water to remove all SDS. After washing, cells were resuspended in 6 mL of 20 mM Tris buffer (pH 8.0). The cells were treated with 800 ug DNase for 24 hrs followed by Trypsin (800 ug) for another 24 hrs (37 ^O^C shaking at 80 rpm). Cells were then boiled for 30 min. Lysozyme (250 ug/mL) was added and cells incubated for 24 hrs at 37 ^O^C shaking at 250 rpm. Following lysozyme digestion, cells were frowzen and lyophilized, then dissolved in 0.375 M sodium borate buffer (pH 9.0) prepared in HPLC grade water. Sodium borohydride (10 mg) in 1 mL borate buffer was added for 30 min, then quenched with 125 uL phosphoric acid. The cell wall digest was filtered using a 0.45 um and 30K centrifuge filter respectively, freezed and lyophilized. The digest was then analyzed by LC-MS.

**Scheme S1. Synthesis of Fl-Tri-(Lys)**.
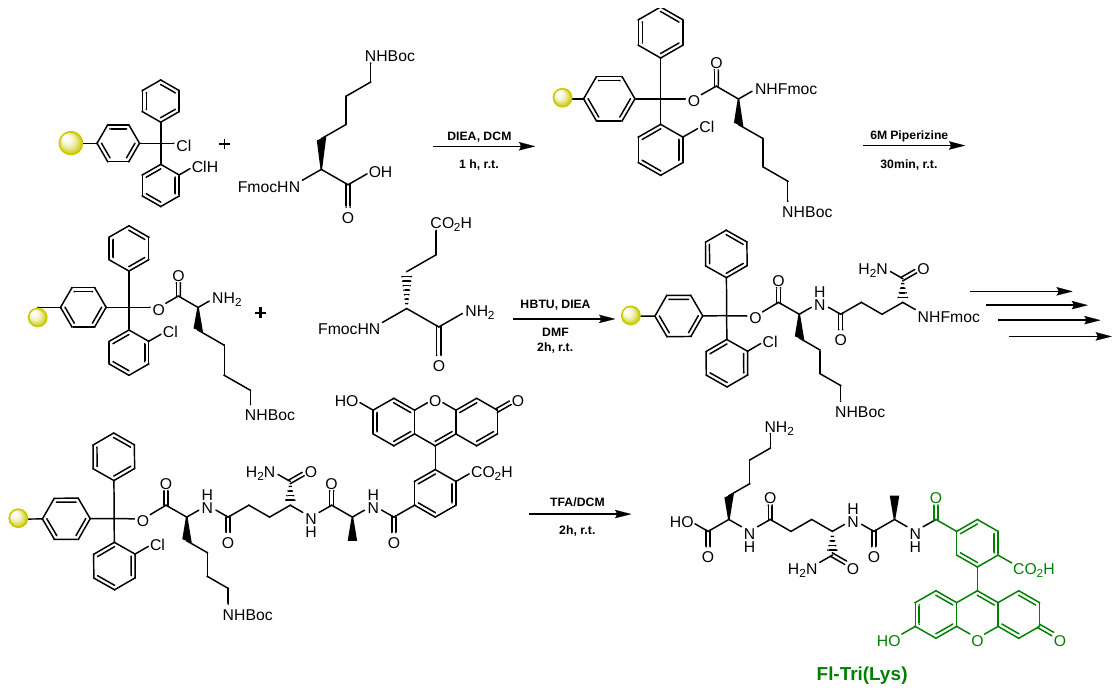

A 25 mL peptide synthesis vessel charged with 2-Chlorotrityl chloride resin (500mg, 0.55mmol) was added Fmoc-L-Lysine(Boc)-OH (1.1 eq, 283 mg, 0.605 mmol) and DIEA (3 eq, 0.286 mL, 1.65 mmol) in dry DCM (15 mL). The resin was agitated for 1 h at ambient temperature and washed with MeOH and DCM (3 x 15 mL each). The Fmoc protecting group was removed with 6 M piperazine/100 mM HOBt in DMF (15 ml) for 30 min at ambient temperature, then washed as before. Fmoc-D-glutamic acid α-amide (3 eq, 607 mg, 1.65 mmol), HBTU (3 eq, 625 mg, 1.65 mmol), and DIEA (6 eq, 0.574 mL, 3.30 mmol) in DMF (15 mL) were added to the reaction flask and agitated for 2 h at ambient temperature. The Fmoc deprotection and coupling procedure was repeated as before using the same equivalencies with Fmoc-L-Alanine-OH. The Fmoc group of L-alanine was deprotected and resin coupled with 5(6)-carboxyfluorescein (2 eq, 413 mg, 1.1 mmol), HBTU (2 eq, 416 mg, 1.1 mmol) and DIEA (6 eq, 0.574 mL, 3.30 mmol) in DMF (15 mL) shaking overnight. The resin was washed as before and added to a solution of TFA/DCM (2:1, 20 mL) with agitation for 2 h at ambient temperature. The resin was filtered and resulting solution concentrated *in vacuo*. The residue was trituated with cold diethyl ether and purified using reverse phase HPLC using H_2_O/MeOH to yield **Fl-Tri(Lys)**. The sample was analyzed for purity using a Shimadzu LC 2020 with a Phenomenex Luna 5µ C18(2) 100Å (30 x 2.00 mm) column; gradient elution with H_2_O/CH_3_CN.

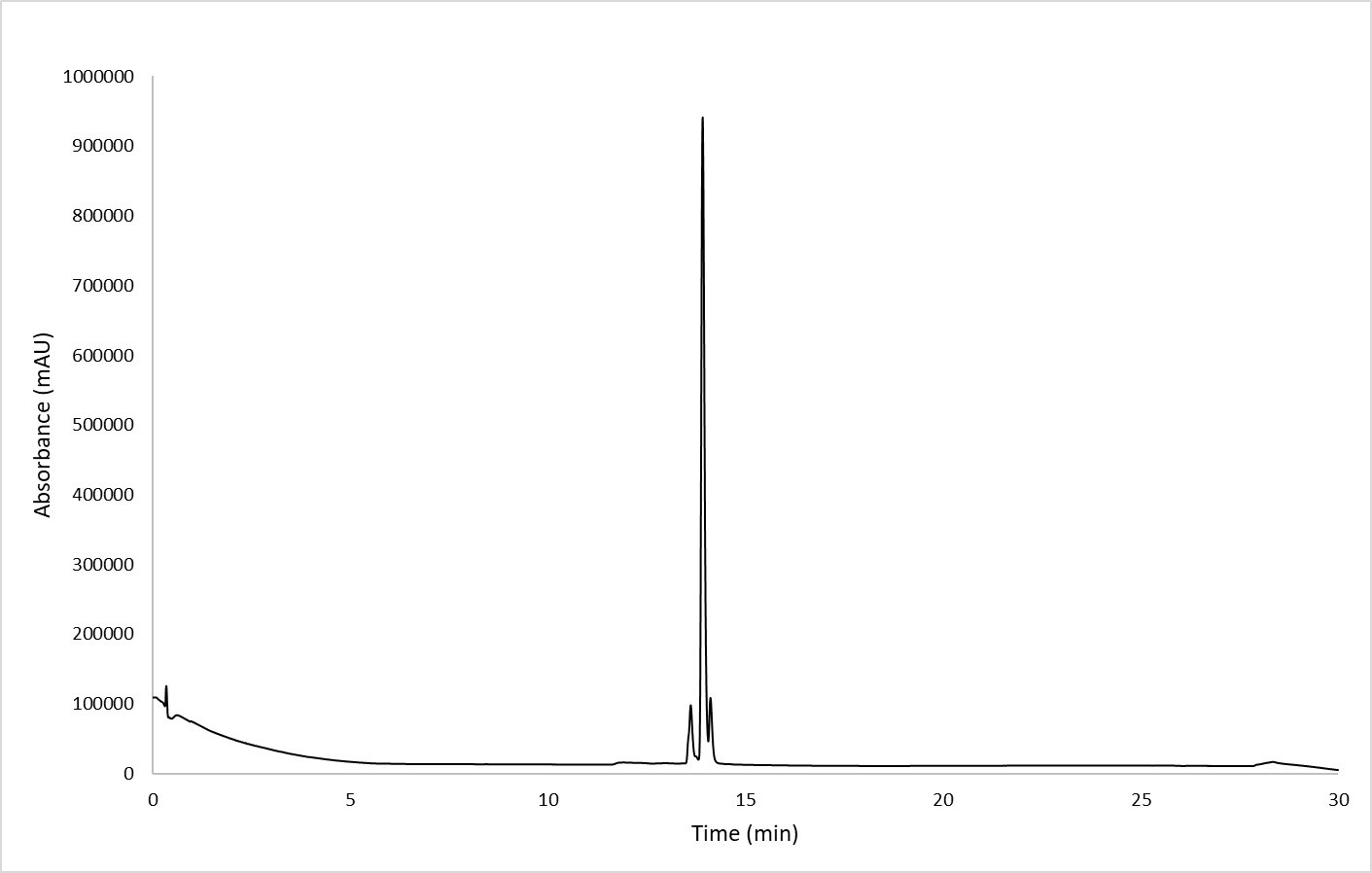

**Scheme S2. Synthesis of Fl-Tri(Asn).**

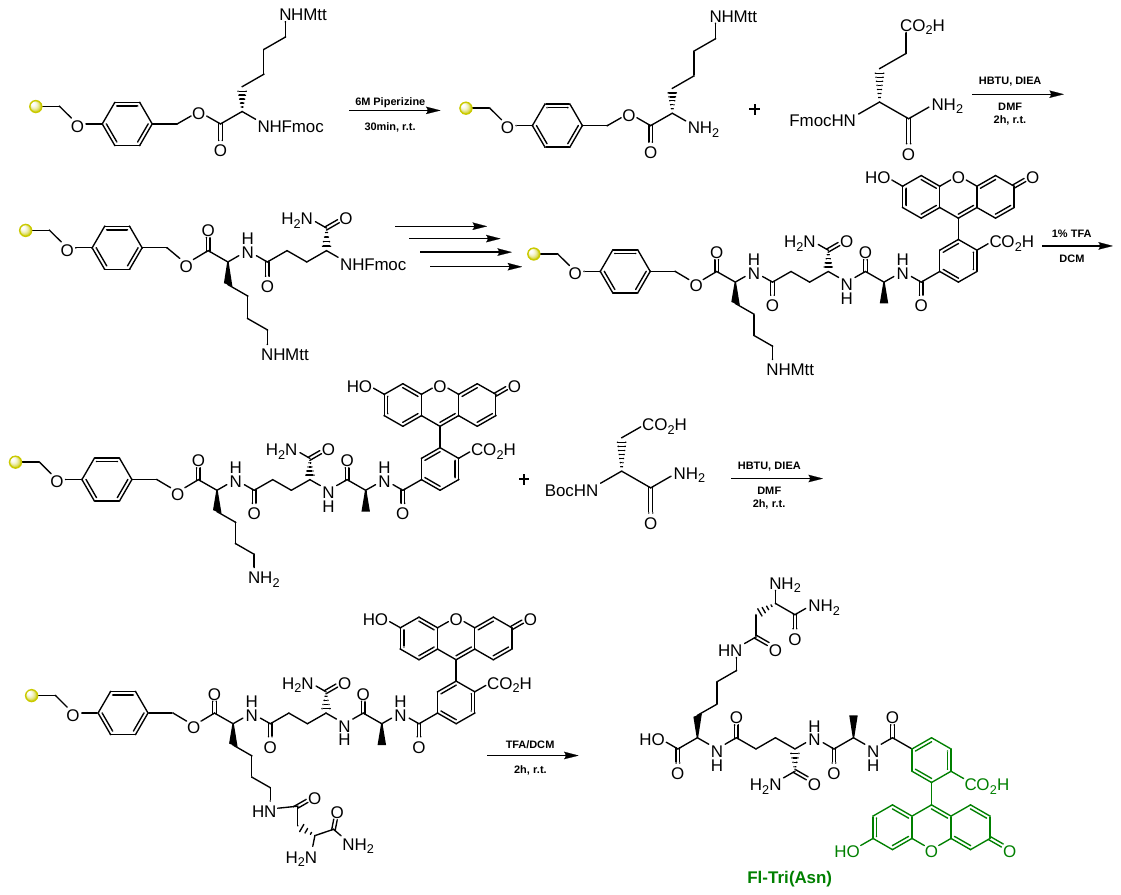

To a 25 mL peptide synthesis vessel charged with Fmoc-Lysine(Mtt)-Wang resin (1.0 g, 0.55mmol). The Fmoc protecting group was removed with 6 M piperazine/100 mM HOBt in DMF (15 ml) for 30 min at ambient temperature, then washed with washed with MeOH and DCM (3 x 15 mL each). Fmoc-D-glutamic acid α-amide (3 eq, 607 mg, 1.65 mmol), HBTU (3 eq, 625 mg, 1.65 mmol), and DIEA (6 eq, 0.574 mL, 3.30 mmol) in DMF (15 mL) were added to the reaction flask and agitated for 2 h at ambient temperature and washed as before. The Fmoc deprotection and coupling procedure was repeated as before using the same equivalencies with Fmoc-L-Alanine-OH. The Fmoc group of L-alanine was deprotected and resin coupled with 5(6)-carboxyfluorescein (2 eq, 413 mg, 1.1 mmol), HBTU (2 eq, 416 mg, 1.1 mmol) and DIEA (6 eq, 0.574 mL, 3.30 mmol) in DMF (15 mL) shaking overnight. The Mtt protecting group was removed by the addition of 1% TFA, 2.5% TIPS, in 10 mL DCM for 15 min, washed and repeated 3 more times. Boc-D-aspartic acid α-amide (3 eq, 382 mg, 1.65 mmol), HBTU (3 eq, 625 mg, 1.65 mmol), and DIEA (6 eq, 0.574 mL, 3.30 mmol) in DMF (15 mL) were added to the reaction flask and agitated for 2 h at ambient temperature and washed as before. A solution of TFA/DCM (2:1, 20 mL) was added to the resin with agitation for 2 h at ambient temperature. The resin was filtered and resulting solution concentrated *in vacuo*. The residue was trituated with cold diethyl ether and purified using reverse phase HPLC using H_2_O/MeOH to yield **Fl-Tri(Asn)**. The sample was analyzed for purity using a Shimadzu LC 2020 with a Phenomenex Luna 5µ C18(2) 100Å (30 x 2.00 mm) column; gradient elution with H_2_O/CH_3_CN.

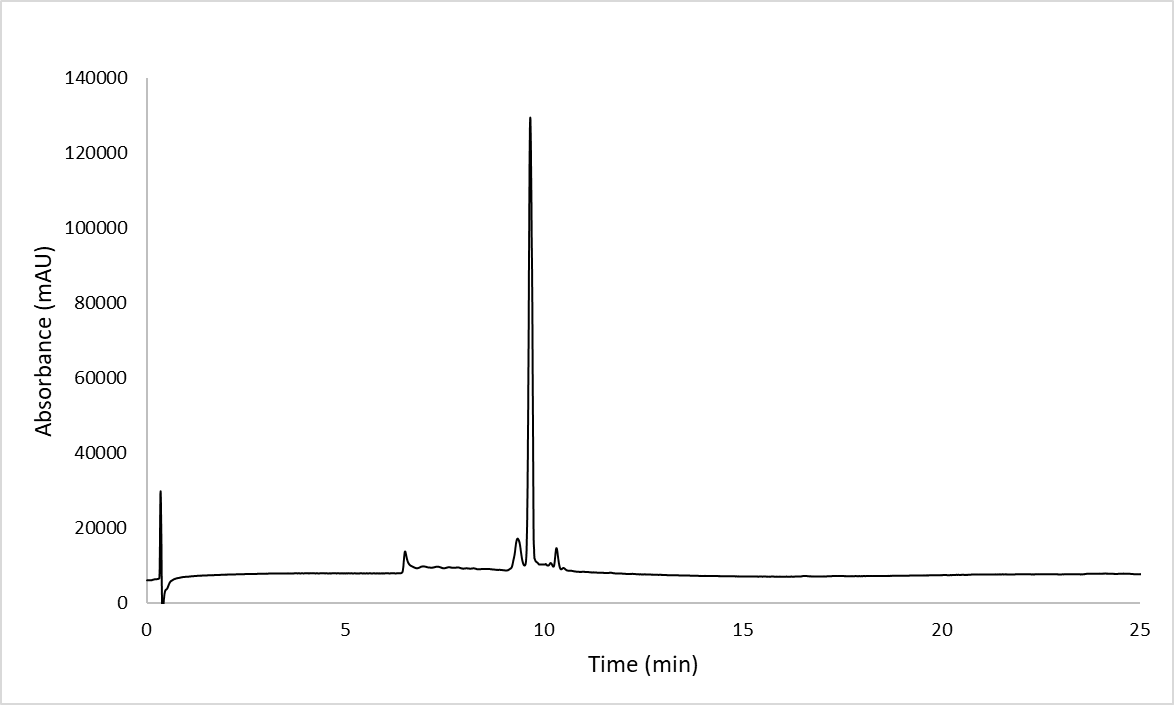

**Scheme S3. Synthesis of Fl-Tri(Asp).**

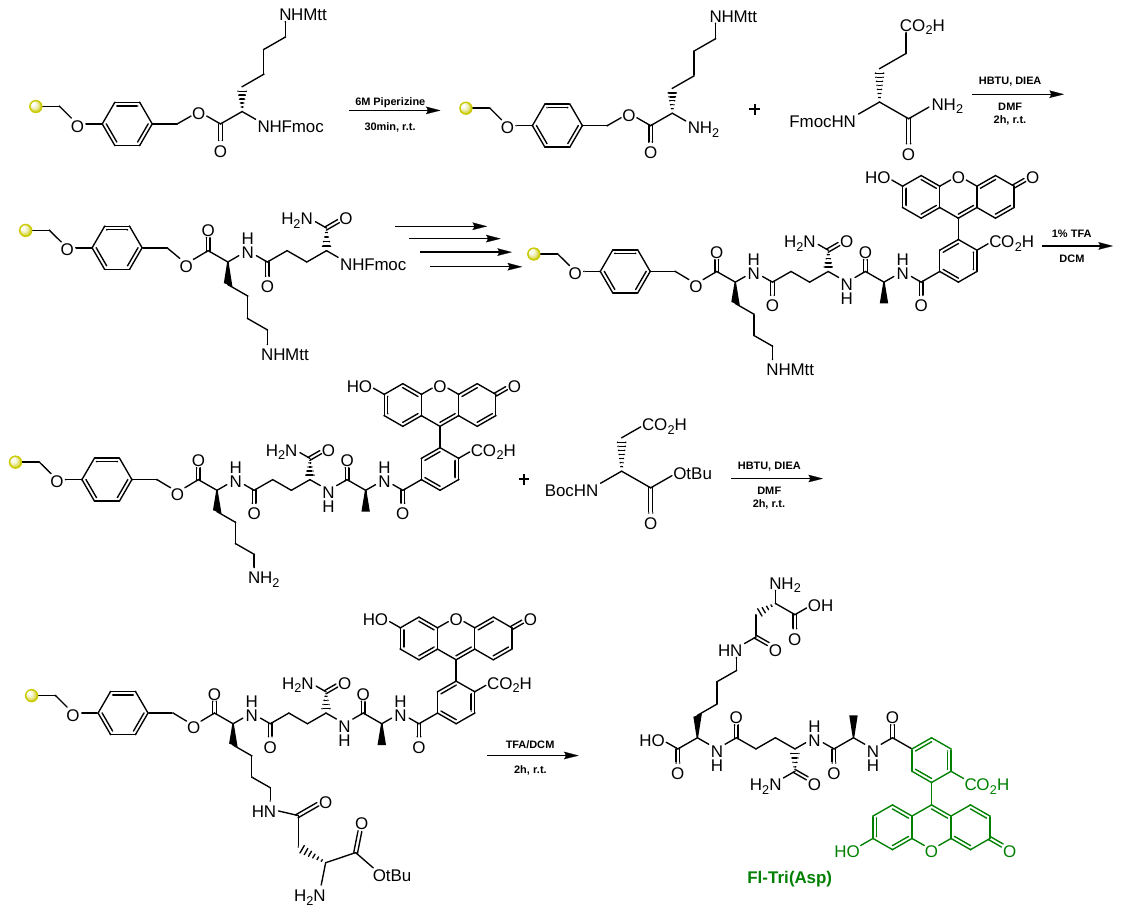

To a 25 mL peptide synthesis vessel charged with Fmoc-Lysine(Mtt)-Wang resin (1.0 g, 0.55mmol). The Fmoc protecting group was removed with 6 M piperazine/100 mM HOBt in DMF (15 ml) for 30 min at ambient temperature, then washed with washed with MeOH and DCM (3 x 15 mL each). Fmoc-D-glutamic acid α-amide (3 eq, 607 mg, 1.65 mmol), HBTU (3 eq, 625 mg, 1.65 mmol), and DIEA (6 eq, 0.574 mL, 3.30 mmol) in DMF (15 mL) were added to the reaction flask and agitated for 2 h at ambient temperature and washed as before. The Fmoc deprotection and coupling procedure was repeated as before using the same equivalencies with Fmoc-L-Alanine-OH. The Fmoc group of L-alanine was deprotected and resin coupled with 5(6)-carboxyfluorescein (2 eq, 413 mg, 1.1 mmol), HBTU (2 eq, 416 mg, 1.1 mmol) and DIEA (6 eq, 0.574 mL, 3.30 mmol) in DMF (15 mL) shaking overnight. The Mtt protecting group was removed by the addition of 1% TFA, 2.5% TIPS, in 10 mL DCM for 15 min, washed and repeated 3 more times. Boc-D-aspartic acid α-*tert*-butyl ester (3 eq, 476 mg, 1.65 mmol), HBTU (3 eq, 625 mg, 1.65 mmol), and DIEA (6 eq, 0.574 mL, 3.30 mmol) in DMF (15 mL) were added to the reaction flask and agitated for 2 h at ambient temperature and washed as before. A solution of TFA/DCM (2:1, 20 mL) was added to the resin with agitation for 2 h at ambient temperature. The resin was filtered and resulting solution concentrated *in vacuo*. The residue was trituated with cold diethyl ether and purified using reverse phase HPLC using H_2_O/MeOH to yield **Fl-Tri(Asp)**. The sample was analyzed for purity using a Shimadzu LC 2020 with a Phenomenex Luna 5µ C18(2) 100Å (30 x 2.00 mm) column; gradient elution with H_2_O/CH_3_CN.

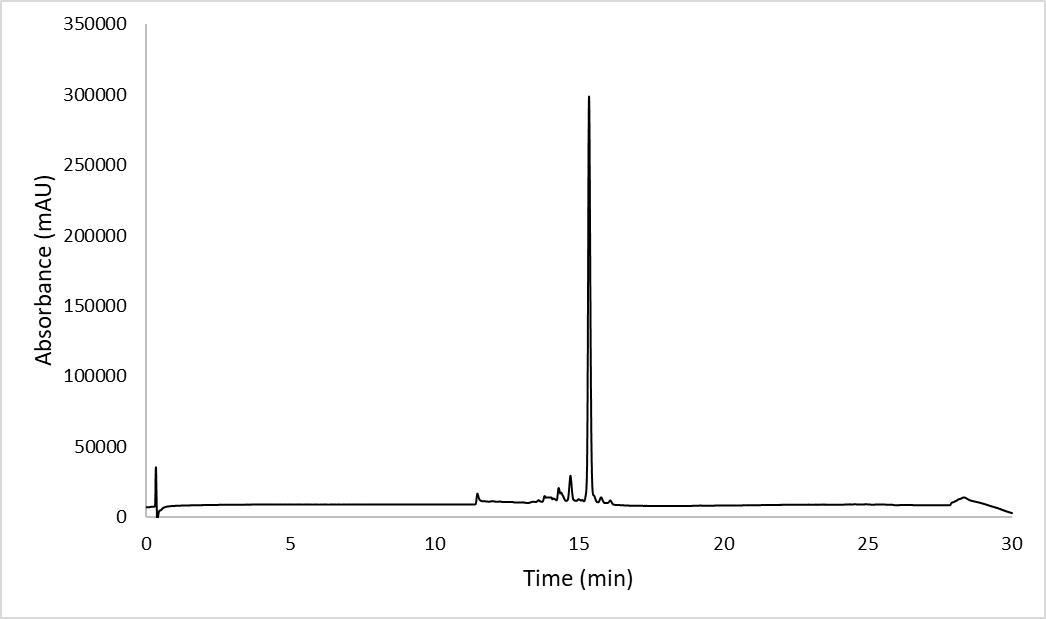

**Scheme S4. Synthesis of Fl-Tri-Glu-(Asn).**

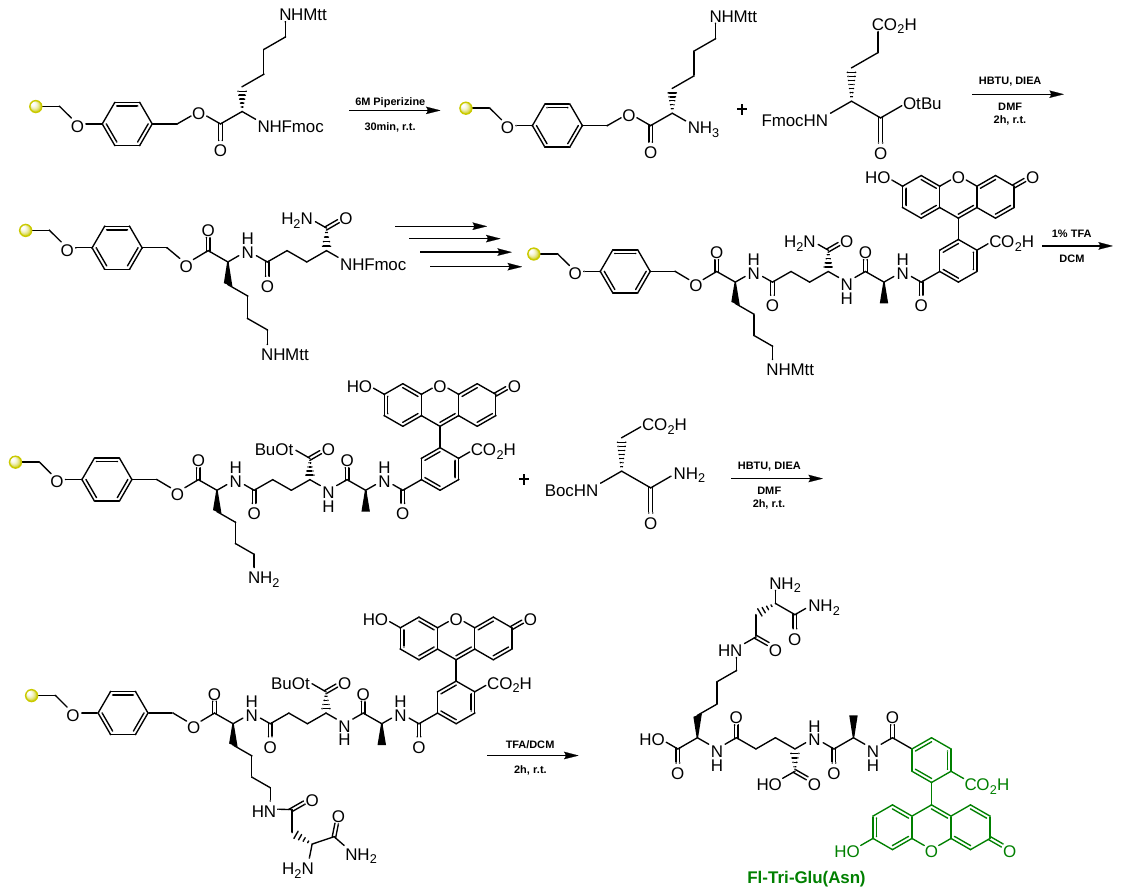

To a 25 mL peptide synthesis vessel charged with Fmoc-Lysine(Mtt)-Wang resin (1.0 g, 0.55mmol). The Fmoc protecting group was removed with 6 M piperazine/100 mM HOBt in DMF (15 ml) for 30 min at ambient temperature, then washed with washed with MeOH and DCM (3 x 15 mL each). Fmoc-D-glutamic acid-a-*tert*-butyl ester (3 eq, 701 mg, 1.65 mmol), HBTU (3 eq, 625 mg, 1.65 mmol), and DIEA (6 eq, 0.574 mL, 3.30 mmol) in DMF (15 mL) were added to the reaction flask and agitated for 2 h at ambient temperature and washed as before. The Fmoc deprotection and coupling procedure was repeated as before using the same equivalencies with Fmoc-L-Alanine-OH. The Fmoc group of L-alanine was deprotected and resin coupled with 5(6)-carboxyfluorescein (2 eq, 413 mg, 1.1 mmol), HBTU (2 eq, 416 mg, 1.1 mmol) and DIEA (6 eq, 0.574 mL, 3.30 mmol) in DMF (15 mL) shaking overnight. The Mtt protecting group was removed by the addition of 1% TFA, 2.5% TIPS, in 10 mL DCM for 15 min, washed and repeated 3 more times. Boc-D-aspartic acid α-amide (3 eq, 382 mg, 1.65 mmol), HBTU (3 eq, 625 mg, 1.65 mmol), and DIEA (6 eq, 0.574 mL, 3.30 mmol) in DMF (15 mL) were added to the reaction flask and agitated for 2 h at ambient temperature and washed as before. A solution of TFA/DCM (2:1, 20 mL) was added to the resin with agitation for 2 h at ambient temperature. The resin was filtered and resulting solution concentrated *in vacuo*. The residue was trituated with cold diethyl ether and purified using reverse phase HPLC using H_2_O/MeOH to yield **Fl-Tri-Glu(Asp)**. The sample was analyzed for purity using a Shimadzu LC 2020 with a Phenomenex Luna 5µ C18(2) 100Å (30 x 2.00 mm) column; gradient elution with H_2_O/CH_3_CN.

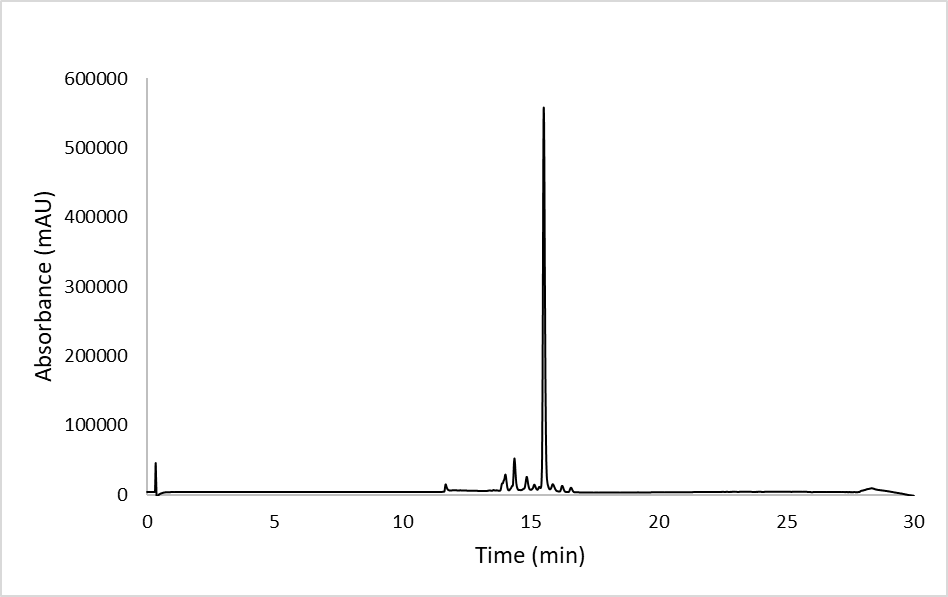

**Scheme S5. Synthesis of Fl-Tri(L-Ala-L-Ala).**

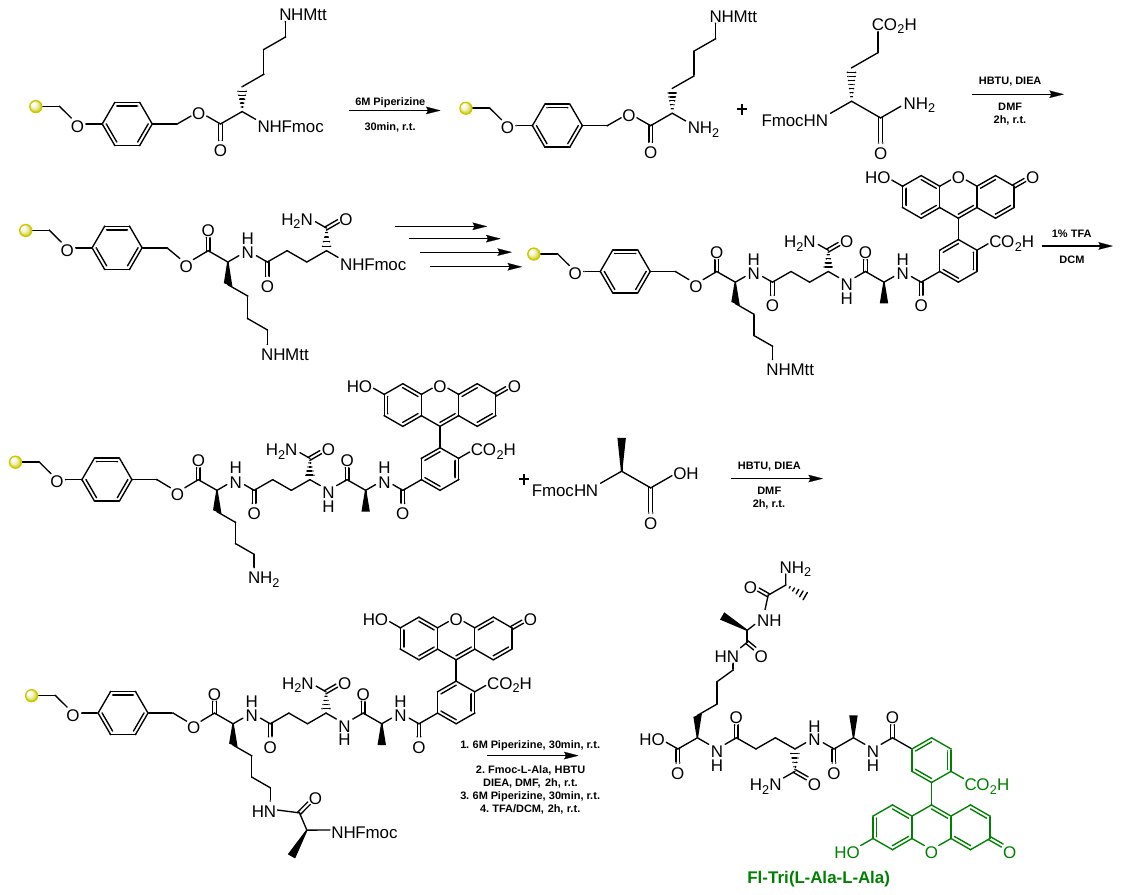

To a 25 mL peptide synthesis vessel charged with Fmoc-Lysine(Mtt)-Wang resin (1.0 g, 0.55mmol). The Fmoc protecting group was removed with 6 M piperazine/100 mM HOBt in DMF (15 ml) for 30 min at ambient temperature, then washed with washed with MeOH and DCM (3 x 15 mL each). Fmoc-D-glutamic acid α-amide (3 eq, 607 mg, 1.65 mmol), HBTU (3 eq, 625 mg, 1.65 mmol), and DIEA (6 eq, 0.574 mL, 3.30 mmol) in DMF (15 mL) were added to the reaction flask and agitated for 2 h at ambient temperature and washed as before. The Fmoc deprotection and coupling procedure was repeated as before using the same equivalencies with Fmoc-L-Alanine-OH. The Fmoc group of L-alanine was deprotected and resin coupled with 5(6)-carboxyfluorescein (2 eq, 413 mg, 1.1 mmol), HBTU (2 eq, 416 mg, 1.1 mmol) and DIEA (6 eq, 0.574 mL, 3.30 mmol) in DMF (15 mL) shaking overnight. The Mtt protecting group was removed by the addition of 1% TFA, 2.5% TIPS, in 10 mL DCM for 15 min, washed and repeated 3 more times. Fmoc-L-alanine (3 eq, 513 mg, 1.65 mmol), HBTU (3 eq, 625 mg, 1.65 mmol), and DIEA (6 eq, 0.574 mL, 3.30 mmol) in DMF (15 mL) were added to the reaction flask and agitated for 2 h at ambient temperature and washed as before. The Fmoc group of L-alanine was deprotected and washed as before. Fmoc-L-alanine (3 eq, 513 mg, 1.65 mmol), HBTU (3 eq, 625 mg, 1.65 mmol), and DIEA (6 eq, 0.574 mL, 3.30 mmol) in DMF (15 mL) were added to the reaction flask and agitated for 2 h at ambient temperature and washed as before. The Fmoc protecting group was removed and a solution of TFA/DCM (2:1, 20 mL) was added to the resin with agitation for 2 h at ambient temperature. The resin was filtered and resulting solution concentrated *in vacuo*. The residue was trituated with cold diethyl ether and purified using reverse phase HPLC using H_2_O/MeOH to yield **Fl-Tri(L-Ala-L-Ala)**. The sample was analyzed for purity using a Shimadzu LC 2020 with a Phenomenex Luna 5µ C18(2) 100Å (30 x 2.00 mm) column; gradient elution with H_2_O/CH_3_CN.

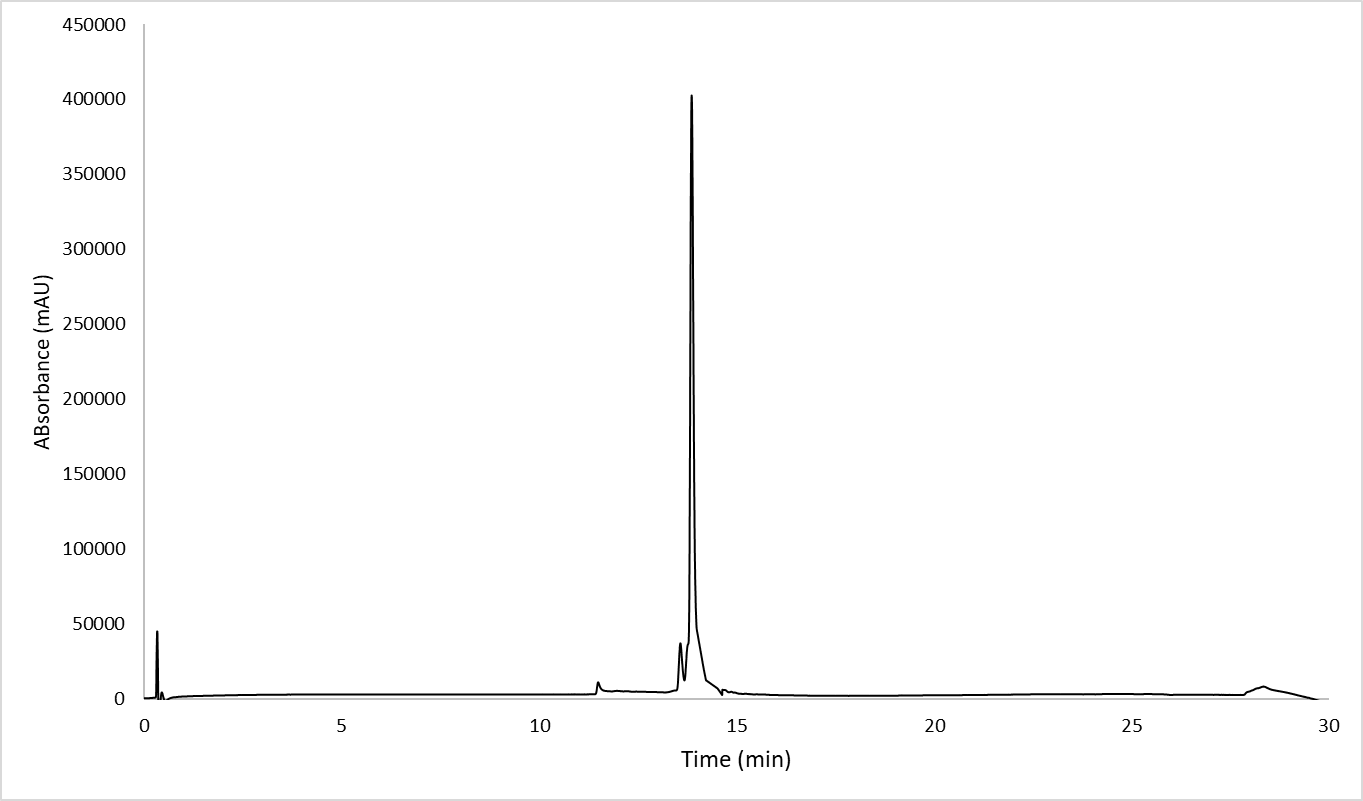

**Scheme S6. Synthesis of Fl-Tri-Glu-(L-Ala-L-Ala).**

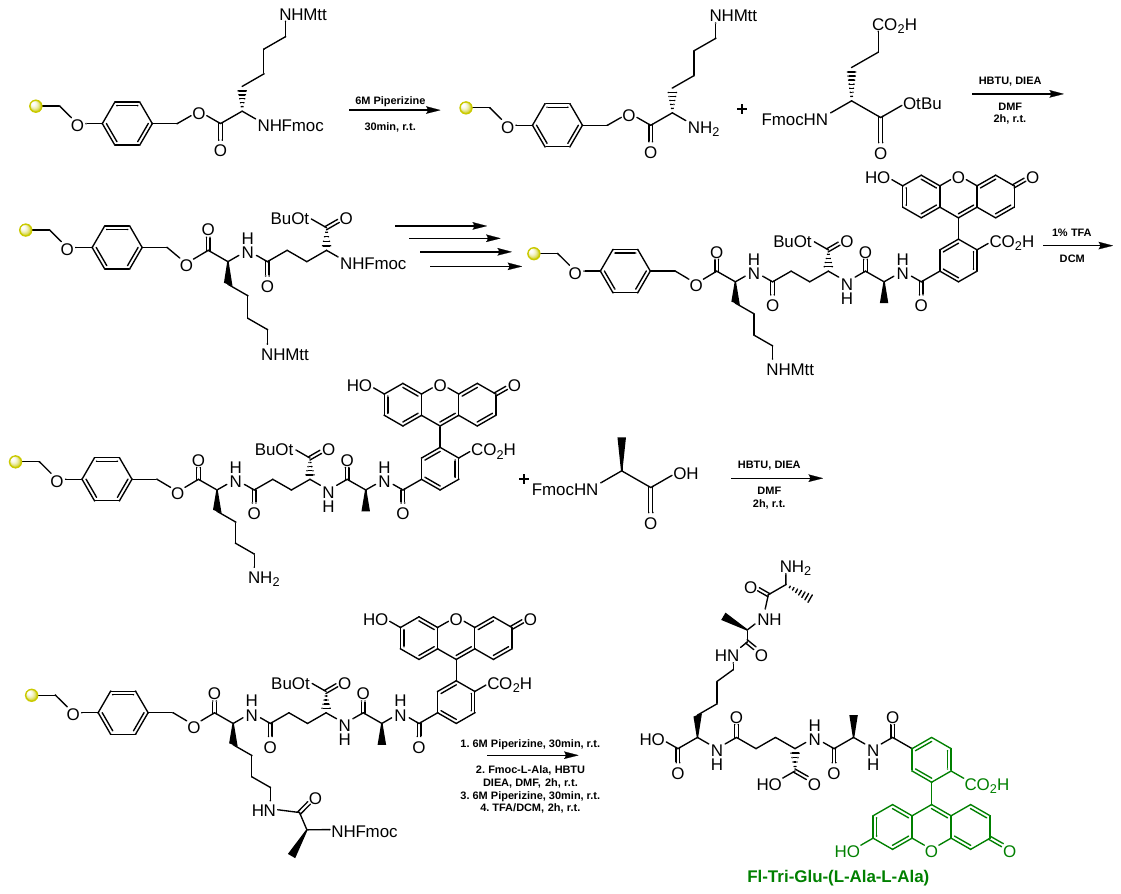

To a 25 mL peptide synthesis vessel charged with Fmoc-Lysine(Mtt)-Wang resin (1.0 g, 0.55mmol). The Fmoc protecting group was removed with 6 M piperazine/100 mM HOBt in DMF (15 ml) for 30 min at ambient temperature, then washed with washed with MeOH and DCM (3 x 15 mL each). Fmoc-D-glutamic acid-a-*tert*-butyl ester (3 eq, 701 mg, 1.65 mmol), HBTU (3 eq, 625 mg, 1.65 mmol), and DIEA (6 eq, 0.574 mL, 3.30 mmol) in DMF (15 mL) were added to the reaction flask and agitated for 2 h at ambient temperature and washed as before. The Fmoc deprotection and coupling procedure was repeated as before using the same equivalencies with Fmoc-L-Alanine-OH. The Fmoc group of L-alanine was deprotected and resin coupled with 5(6)-carboxyfluorescein (2 eq, 413 mg, 1.1 mmol), HBTU (2 eq, 416 mg, 1.1 mmol) and DIEA (6 eq, 0.574 mL, 3.30 mmol) in DMF (15 mL) shaking overnight. The Mtt protecting group was removed by the addition of 1% TFA, 2.5% TIPS, in 10 mL DCM for 15 min, washed and repeated 3 more times. Fmoc-L-alanine (3 eq, 513 mg, 1.65 mmol), HBTU (3 eq, 625 mg, 1.65 mmol), and DIEA (6 eq, 0.574 mL, 3.30 mmol) in DMF (15 mL) were added to the reaction flask and agitated for 2 h at ambient temperature and washed as before. The Fmoc group of L-alanine was deprotected and washed as before. Fmoc-L-alanine (3 eq, 513 mg, 1.65 mmol), HBTU (3 eq, 625 mg, 1.65 mmol), and DIEA (6 eq, 0.574 mL, 3.30 mmol) in DMF (15 mL) were added to the reaction flask and agitated for 2 h at ambient temperature and washed as before. The Fmoc protecting group was removed and a solution of TFA/DCM (2:1, 20 mL) was added to the resin with agitation for 2 h at ambient temperature. The resin was filtered and resulting solution concentrated *in vacuo*. The residue was trituated with cold diethyl ether and purified using reverse phase HPLC using H_2_O/MeOH to yield **Fl-Tri-Glu(L-Ala-L-Ala)**. The sample was analyzed for purity using a Shimadzu LC 2020 with a Phenomenex Luna 5µ C18(2) 100Å (30 x 2.00 mm) column; gradient elution with H_2_O/CH_3_CN.

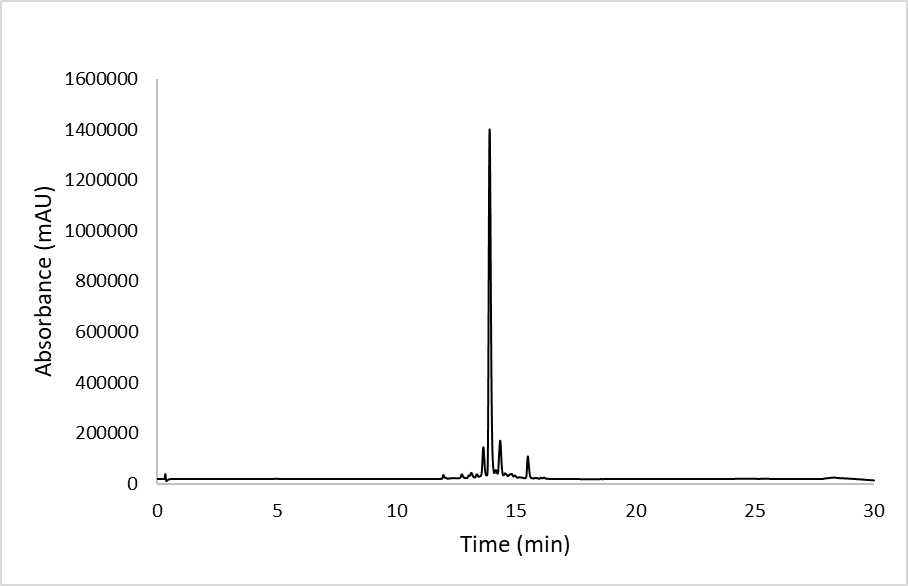

**Scheme S7. Synthesis of Fl-Tri(D-Ala-D-Ala).**

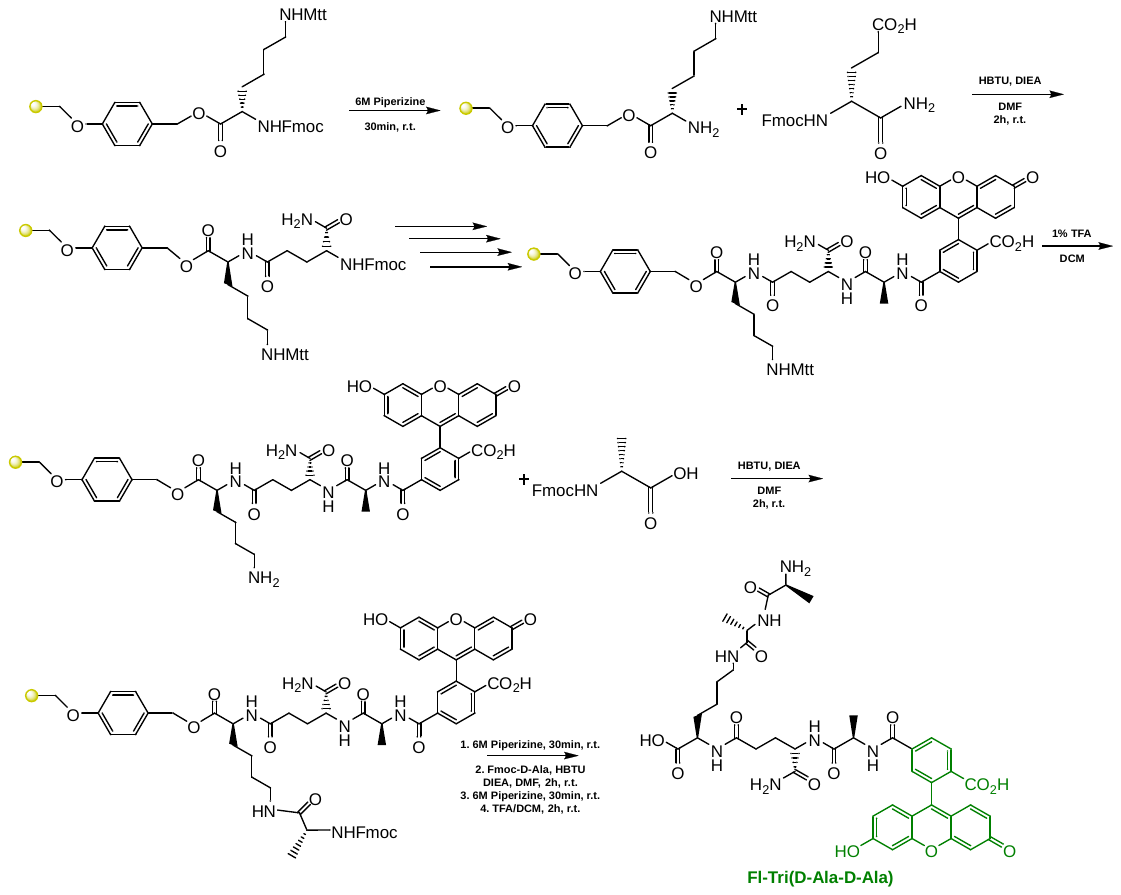

To a 25 mL peptide synthesis vessel charged with Fmoc-Lysine(Mtt)-Wang resin (1.0 g, 0.55mmol). The Fmoc protecting group was removed with 6 M piperazine/100 mM HOBt in DMF (15 ml) for 30 min at ambient temperature, then washed with washed with MeOH and DCM (3 x 15 mL each). Fmoc-D-glutamic acid α-amide (3 eq, 607 mg, 1.65 mmol), HBTU (3 eq, 625 mg, 1.65 mmol), and DIEA (6 eq, 0.574 mL, 3.30 mmol) in DMF (15 mL) were added to the reaction flask and agitated for 2 h at ambient temperature and washed as before. The Fmoc deprotection and coupling procedure was repeated as before using the same equivalencies with Fmoc-L-Alanine-OH. The Fmoc group of L-alanine was deprotected and resin coupled with 5(6)-carboxyfluorescein (2 eq, 413 mg, 1.1 mmol), HBTU (2 eq, 416 mg, 1.1 mmol) and DIEA (6 eq, 0.574 mL, 3.30 mmol) in DMF (15 mL) shaking overnight. The Mtt protecting group was removed by the addition of 1% TFA, 2.5% TIPS, in 10 mL DCM for 15 min, washed and repeated 3 more times. Fmoc-D-alanine (3 eq, 513 mg, 1.65 mmol), HBTU (3 eq, 625 mg, 1.65 mmol), and DIEA (6 eq, 0.574 mL, 3.30 mmol) in DMF (15 mL) were added to the reaction flask and agitated for 2 h at ambient temperature and washed as before. The Fmoc group of D-alanine was deprotected and washed as before. Fmoc-D-alanine (3 eq, 513 mg, 1.65 mmol), HBTU (3 eq, 625 mg, 1.65 mmol), and DIEA (6 eq, 0.574 mL, 3.30 mmol) in DMF (15 mL) were added to the reaction flask and agitated for 2 h at ambient temperature and washed as before. The Fmoc protecting group was removed and a solution of TFA/DCM (2:1, 20 mL) was added to the resin with agitation for 2 h at ambient temperature. The resin was filtered and resulting solution concentrated *in vacuo*. The residue was trituated with cold diethyl ether and purified using reverse phase HPLC using H_2_O/MeOH to yield **Fl-Tri(D-Ala-D-Ala)**. The sample was analyzed for purity using a Shimadzu LC 2020 with a Phenomenex Luna 5µ C18(2) 100Å (30 x 2.00 mm) column; gradient elution with H_2_O/CH_3_CN.

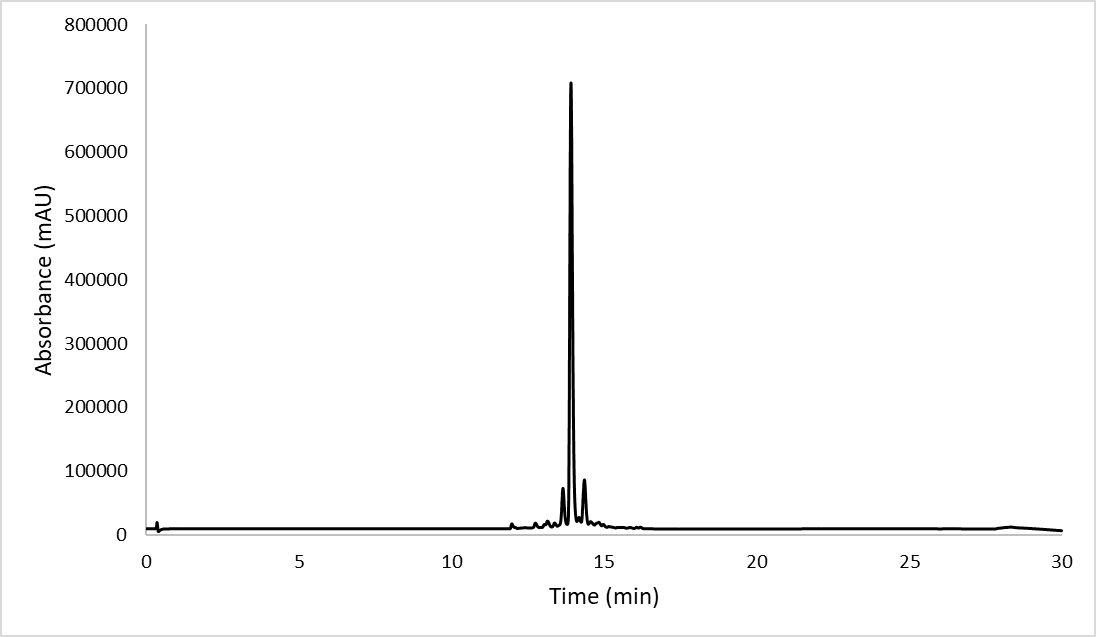

**Scheme S8. Synthesis of Fl-Tri(L-Ala-L-Ala-Ac).**

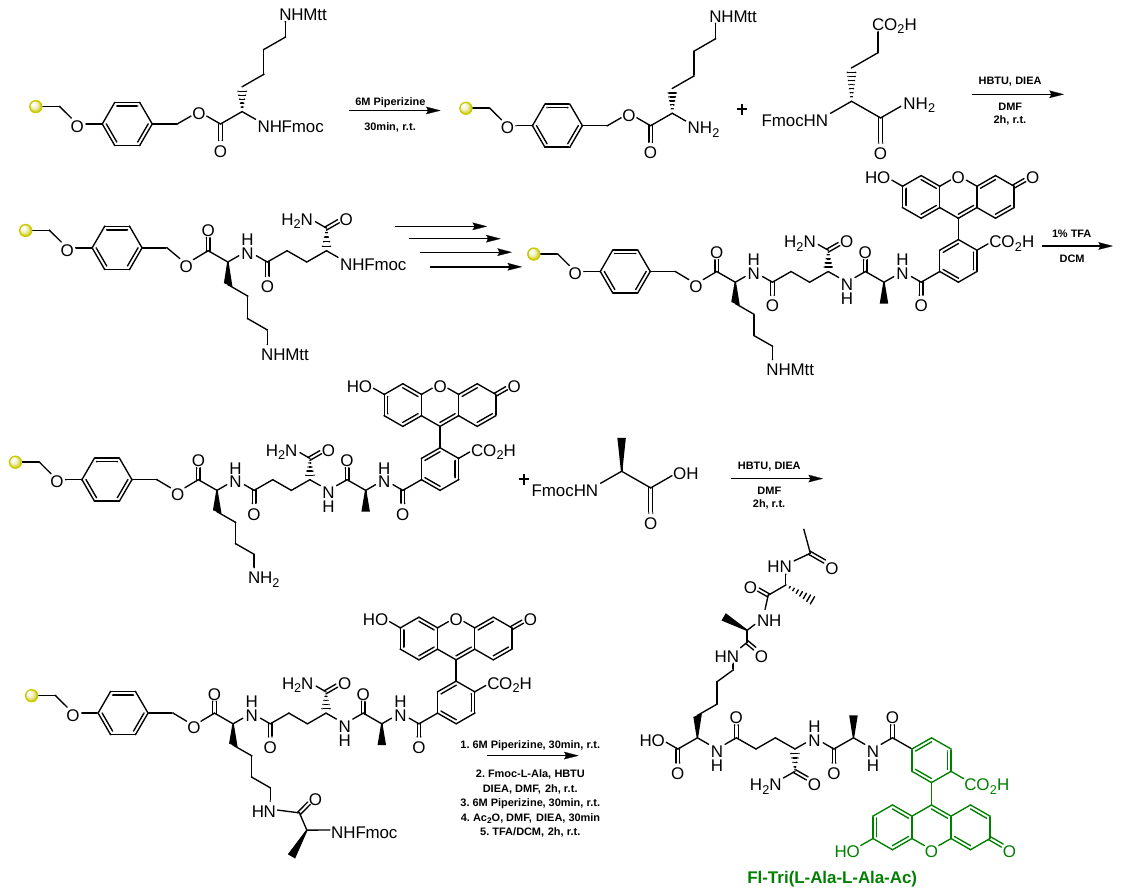

To a 25 mL peptide synthesis vessel charged with Fmoc-Lysine(Mtt)-Wang resin (1.0 g, 0.55mmol). The Fmoc protecting group was removed with 6 M piperazine/100 mM HOBt in DMF (15 ml) for 30 min at ambient temperature, then washed with washed with MeOH and DCM (3 x 15 mL each). Fmoc-D-glutamic acid α-amide (3 eq, 607 mg, 1.65 mmol), HBTU (3 eq, 625 mg, 1.65 mmol), and DIEA (6 eq, 0.574 mL, 3.30 mmol) in DMF (15 mL) were added to the reaction flask and agitated for 2 h at ambient temperature and washed as before. The Fmoc deprotection and coupling procedure was repeated as before using the same equivalencies with Fmoc-L-Alanine-OH. The Fmoc group of L-alanine was deprotected and resin coupled with 5(6)-carboxyfluorescein (2 eq, 413 mg, 1.1 mmol), HBTU (2 eq, 416 mg, 1.1 mmol) and DIEA (6 eq, 0.574 mL, 3.30 mmol) in DMF (15 mL) shaking overnight. The Mtt protecting group was removed by the addition of 1% TFA, 2.5% TIPS, in 10 mL DCM for 15 min, washed and repeated 3 more times. Fmoc-L-alanine (3 eq, 513 mg, 1.65 mmol), HBTU (3 eq, 625 mg, 1.65 mmol), and DIEA (6 eq, 0.574 mL, 3.30 mmol) in DMF (15 mL) were added to the reaction flask and agitated for 2 h at ambient temperature and washed as before. The Fmoc group of L-alanine was deprotected and washed as before. Fmoc-L-alanine (3 eq, 513 mg, 1.65 mmol), HBTU (3 eq, 625 mg, 1.65 mmol), and DIEA (6 eq, 0.574 mL, 3.30 mmol) in DMF (15 mL) were added to the reaction flask and agitated for 2 h at ambient temperature and washed as before. The Fmoc protecting group was removed and a solution of acetic anhydride (5 eq, 0.260 mL) and DIEA (10 eq, 0.956 mL) in DMF was added and resin shaken for 30 min at ambient temperature. The resin was washed and added to a solution of TFA/DCM (2:1, 20 mL) with agitation for 2 h at ambient temperature. The resin was filtered and resulting solution concentrated *in vacuo*. The residue was trituated with cold diethyl ether and purified using reverse phase HPLC using H_2_O/MeOH to yield **Fl-Tri(L-Ala-L-Ala-Ac).** The sample was analyzed for purity using a Shimadzu LC 2020 with a Phenomenex Luna 5µ C18(2) 100Å (30 x 2.00 mm) column; gradient elution with H_2_O/CH_3_CN.

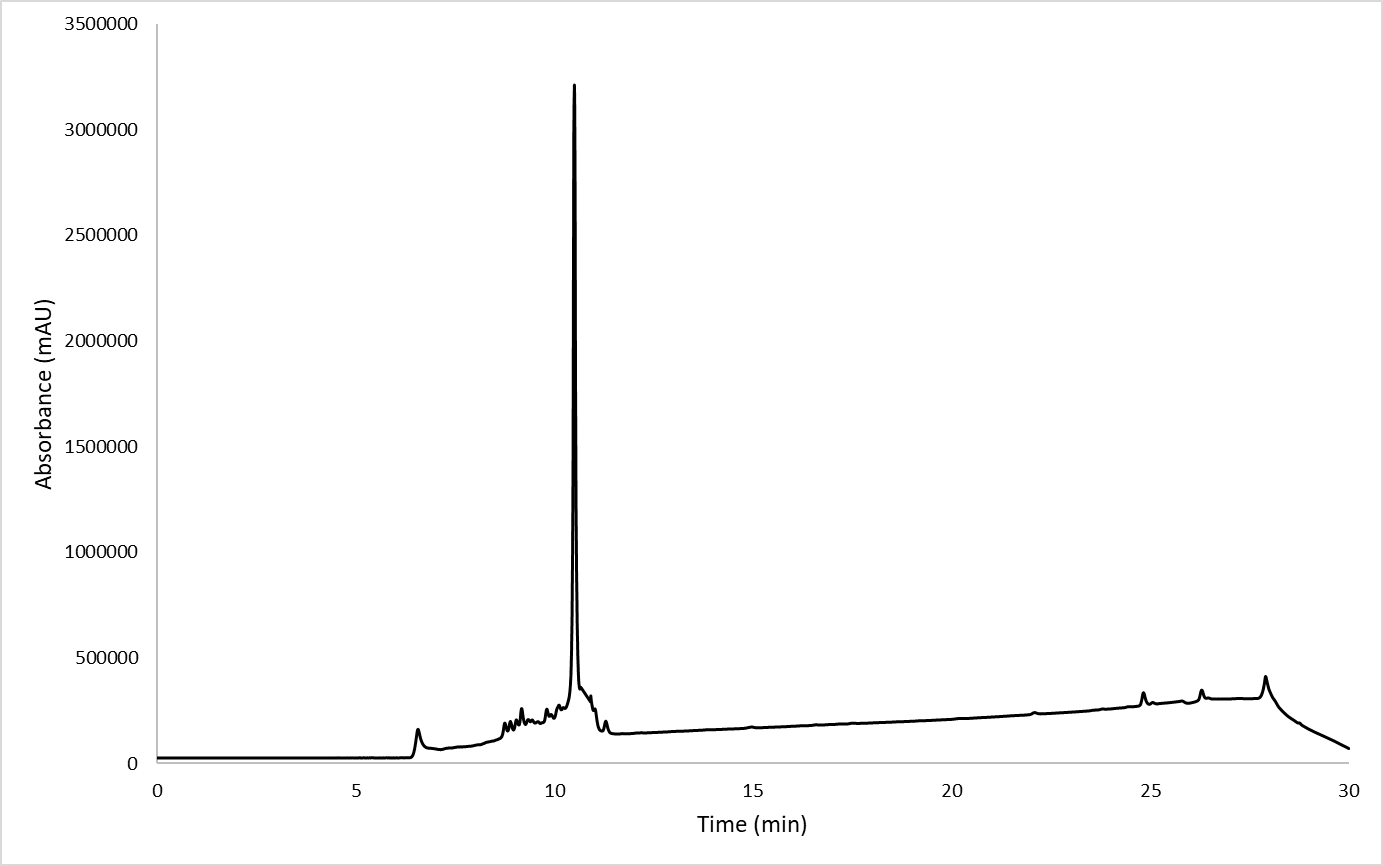

**Scheme S9. Synthesis of Fl-Tri(Gly_1_).**

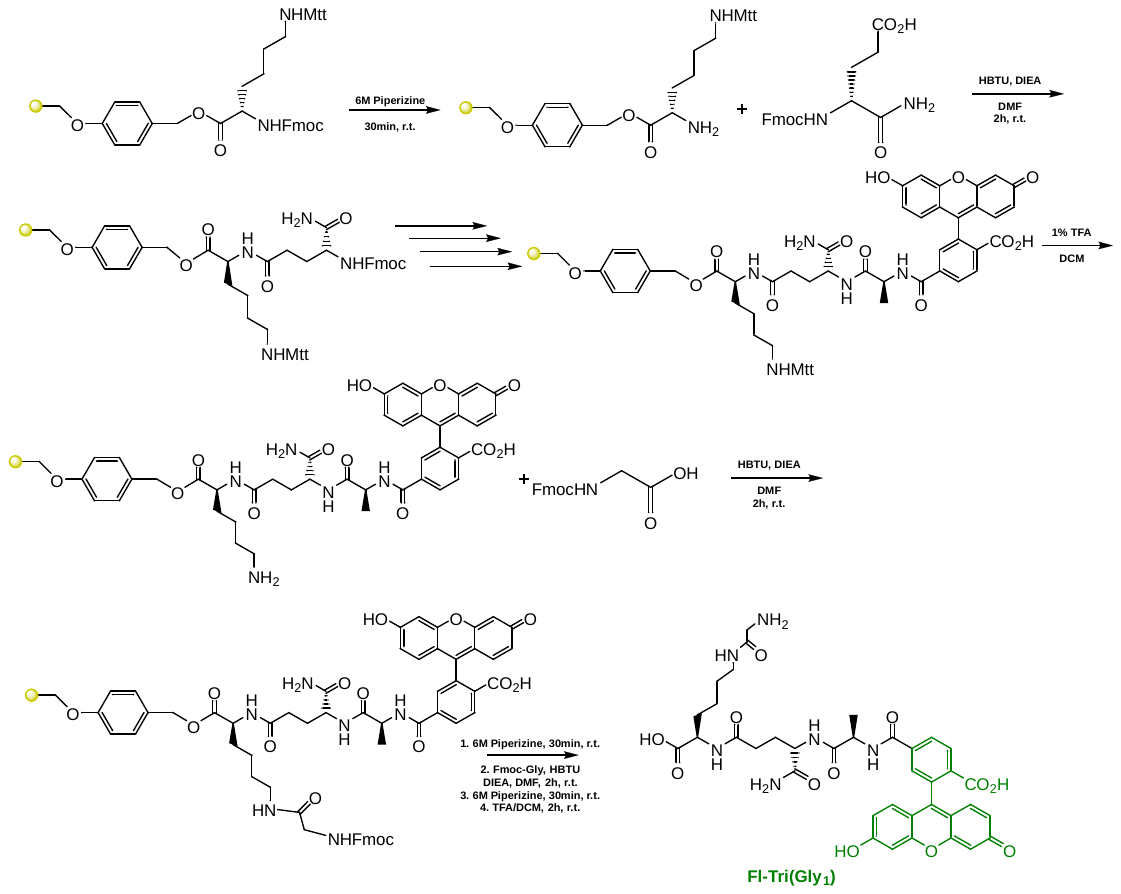

To a 25 mL peptide synthesis vessel charged with Fmoc-Lysine(Mtt)-Wang resin (1.0 g, 0.55mmol). The Fmoc protecting group was removed with 6 M piperazine/100 mM HOBt in DMF (15 ml) for 30 min at ambient temperature, then washed with washed with MeOH and DCM (3 x 15 mL each). Fmoc-D-glutamic acid α-amide (3 eq, 607 mg, 1.65 mmol), HBTU (3 eq, 625 mg, 1.65 mmol), and DIEA (6 eq, 0.574 mL, 3.30 mmol) in DMF (15 mL) were added to the reaction flask and agitated for 2 h at ambient temperature and washed as before. The Fmoc deprotection and coupling procedure was repeated as before using the same equivalencies with Fmoc-L-Alanine-OH. The Fmoc group of L-alanine was deprotected and resin coupled with 5(6)-carboxyfluorescein (2 eq, 413 mg, 1.1 mmol), HBTU (2 eq, 416 mg, 1.1 mmol) and DIEA (6 eq, 0.574 mL, 3.30 mmol) in DMF (15 mL) shaking overnight. The Mtt protecting group was removed by the addition of 1% TFA, 2.5% TIPS, in 10 mL DCM for 15 min, washed and repeated 3 more times. Fmoc-glycine (3 eq, 513 mg, 1.65 mmol), HBTU (3 eq, 625 mg, 1.65 mmol), and DIEA (6 eq, 0.574 mL, 3.30 mmol) in DMF (15 mL) were added to the reaction flask and agitated for 2 h at ambient temperature and washed as before. The Fmoc group of glycine was deprotected, resin washed, and a solution of TFA/DCM (2:1, 20 mL) was added with agitation for 2 h at ambient temperature. The resin was filtered and resulting solution concentrated *in vacuo*. The residue was trituated with cold diethyl ether and purified using reverse phase HPLC using H_2_O/MeOH to yield **Fl-Tri(Gly_1_)**. The sample was analyzed for purity using a Shimadzu LC 2020 with a Phenomenex Luna 5µ C18(2) 100Å (30 x 2.00 mm) column; gradient elution with H_2_O/CH_3_CN.

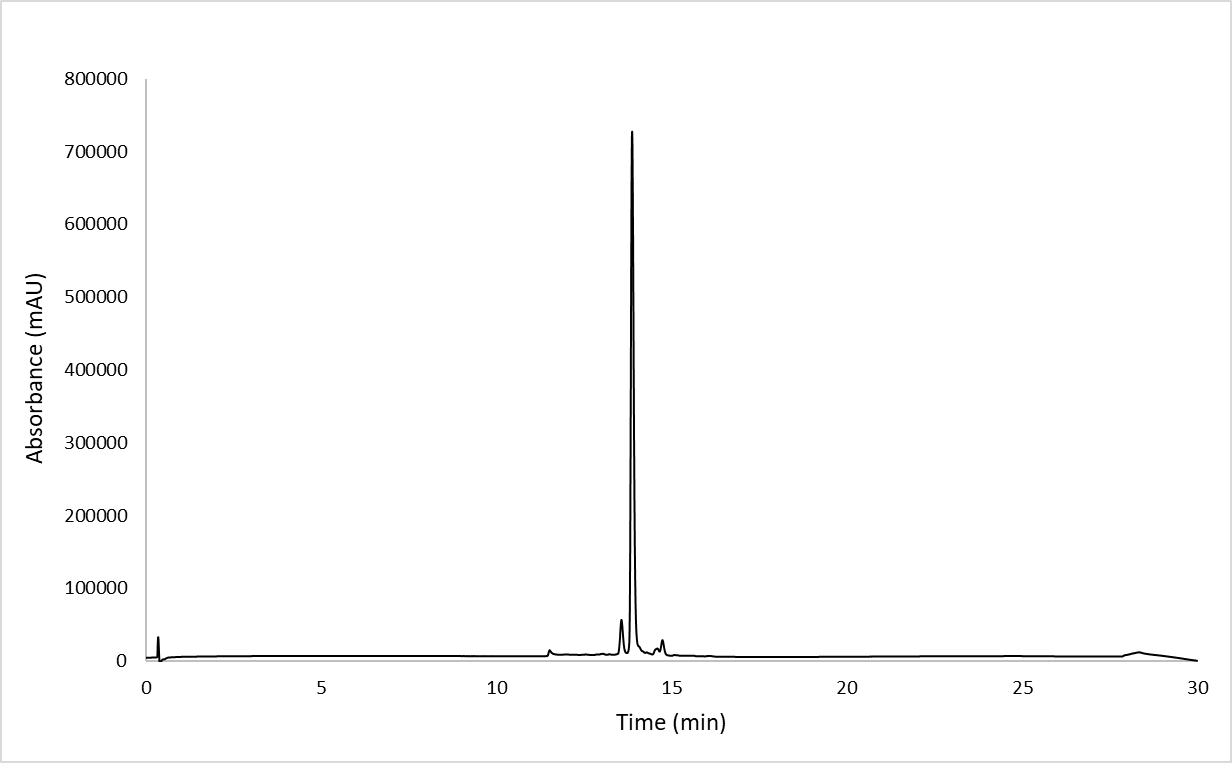

**Scheme S10. Synthesis of Fl-Tri(Gly_2_).**

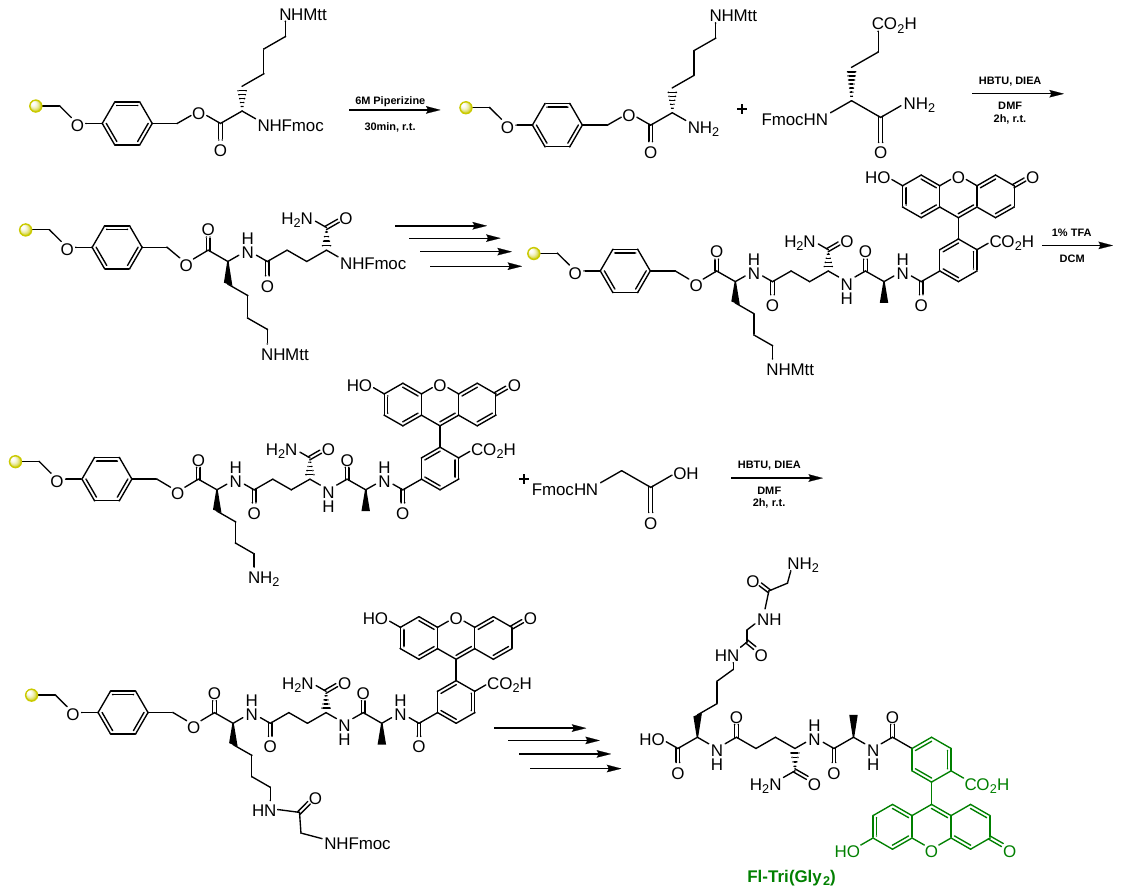

To a 25 mL peptide synthesis vessel charged with Fmoc-Lysine(Mtt)-Wang resin (1.0 g, 0.55mmol). The Fmoc protecting group was removed with 6 M piperazine/100 mM HOBt in DMF (15 ml) for 30 min at ambient temperature, then washed with washed with MeOH and DCM (3 x 15 mL each). Fmoc-D-glutamic acid α-amide (3 eq, 607 mg, 1.65 mmol), HBTU (3 eq, 625 mg, 1.65 mmol), and DIEA (6 eq, 0.574 mL, 3.30 mmol) in DMF (15 mL) were added to the reaction flask and agitated for 2 h at ambient temperature and washed as before. The Fmoc deprotection and coupling procedure was repeated as before using the same equivalencies with Fmoc-L-Alanine-OH. The Fmoc group of L-alanine was deprotected and resin coupled with 5(6)-carboxyfluorescein (2 eq, 413 mg, 1.1 mmol), HBTU (2 eq, 416 mg, 1.1 mmol) and DIEA (6 eq, 0.574 mL, 3.30 mmol) in DMF (15 mL) shaking overnight. The Mtt protecting group was removed by the addition of 1% TFA, 2.5% TIPS, in 10 mL DCM for 15 min, washed and repeated 3 more times. Fmoc-glycine (3 eq, 513 mg, 1.65 mmol), HBTU (3 eq, 625 mg, 1.65 mmol), and DIEA (6 eq, 0.574 mL, 3.30 mmol) in DMF (15 mL) were added to the reaction flask and agitated for 2 h at ambient temperature and washed as before. The Fmoc group of glycine was deprotected, resin washed, and the Fmoc-glycine procedure was repeated once more. After Fmoc deprotection, a solution of TFA/DCM (2:1, 20 mL) was added to the resin with agitation for 2 h at ambient temperature. The resin was filtered and resulting solution concentrated *in vacuo*. The residue was trituated with cold diethyl ether and purified using reverse phase HPLC using H_2_O/MeOH to yield **Fl-Tri(Gly_2_)**. The sample was analyzed for purity using a Shimadzu LC 2020 with a Phenomenex Luna 5µ C18(2) 100Å (30 x 2.00 mm) column; gradient elution with H_2_O/CH_3_CN.

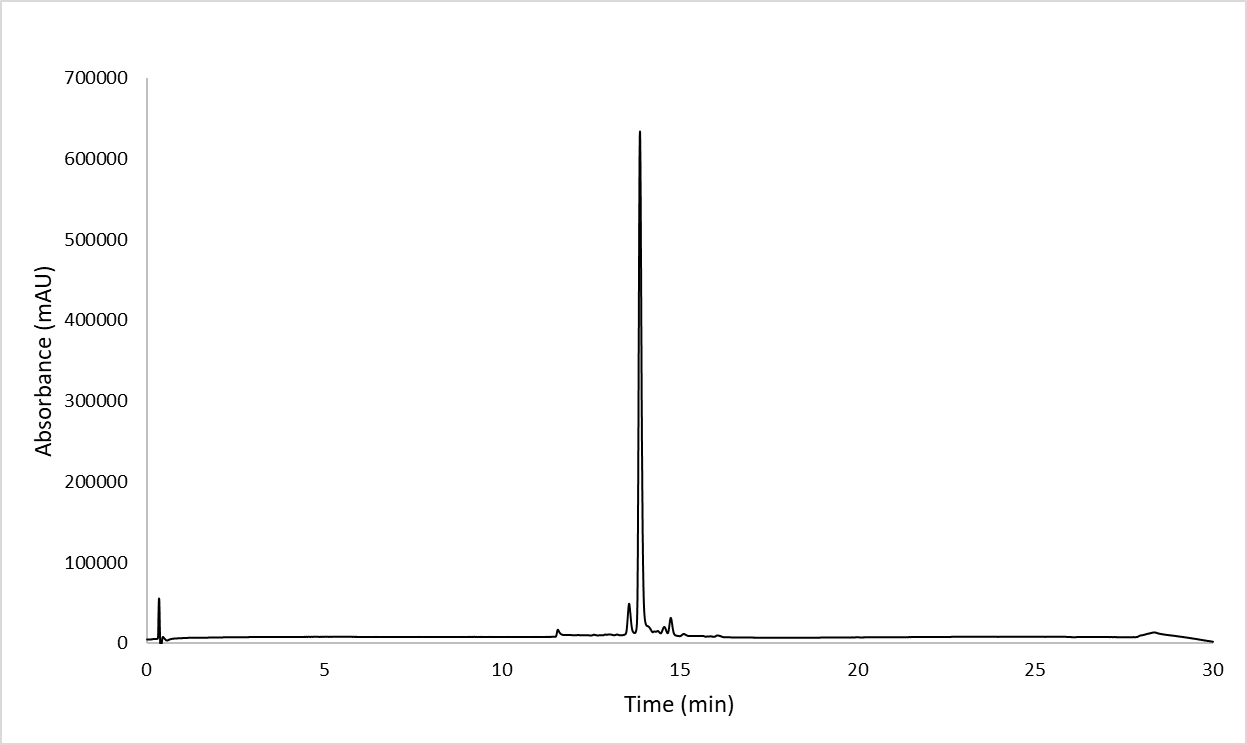

**Scheme S11. Synthesis of Fl-Tri(Gly_3_).**

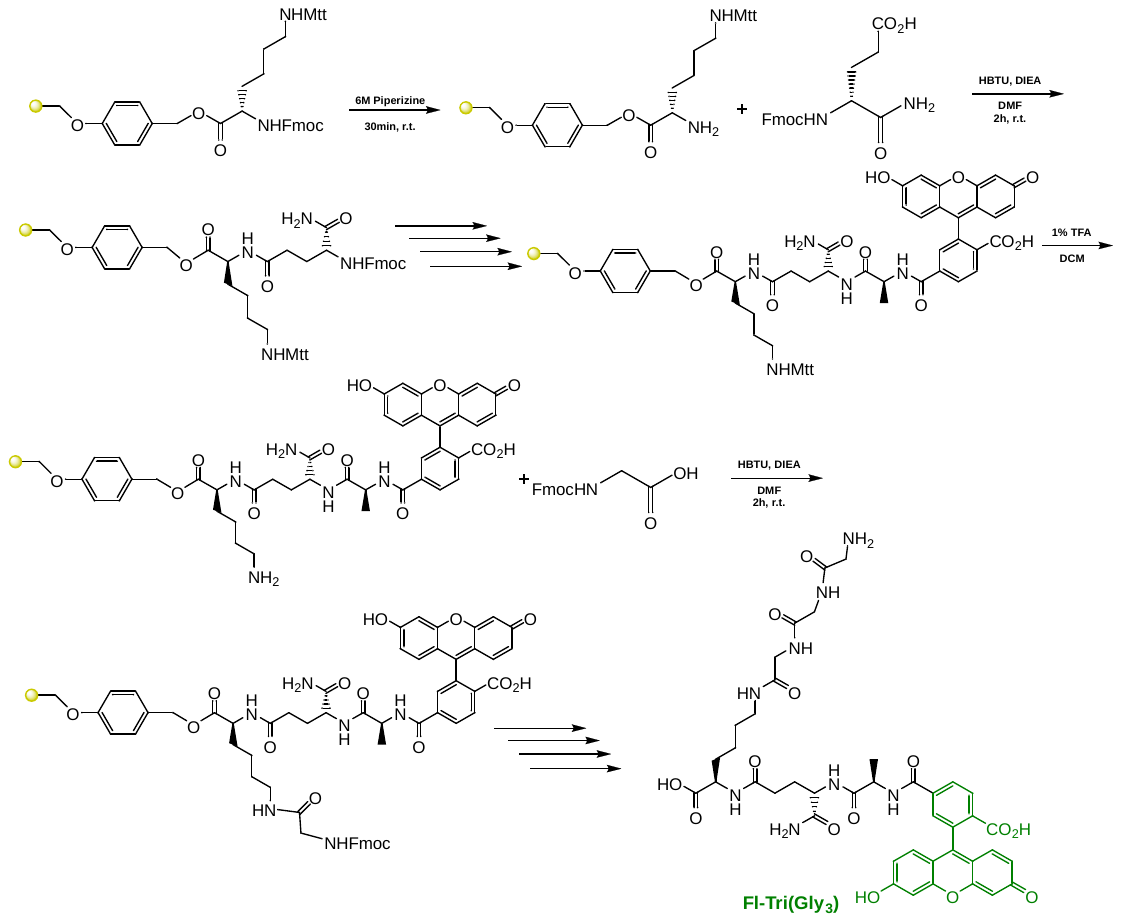

To a 25 mL peptide synthesis vessel charged with Fmoc-Lysine(Mtt)-Wang resin (1.0 g, 0.55mmol). The Fmoc protecting group was removed with 6 M piperazine/100 mM HOBt in DMF (15 ml) for 30 min at ambient temperature, then washed with washed with MeOH and DCM (3 x 15 mL each). Fmoc-D-glutamic acid α-amide (3 eq, 607 mg, 1.65 mmol), HBTU (3 eq, 625 mg, 1.65 mmol), and DIEA (6 eq, 0.574 mL, 3.30 mmol) in DMF (15 mL) were added to the reaction flask and agitated for 2 h at ambient temperature and washed as before. The Fmoc deprotection and coupling procedure was repeated as before using the same equivalencies with Fmoc-L-Alanine-OH. The Fmoc group of L-alanine was deprotected and resin coupled with 5(6)-carboxyfluorescein (2 eq, 413 mg, 1.1 mmol), HBTU (2 eq, 416 mg, 1.1 mmol) and DIEA (6 eq, 0.574 mL, 3.30 mmol) in DMF (15 mL) shaking overnight. The Mtt protecting group was removed by the addition of 1% TFA, 2.5% TIPS, in 10 mL DCM for 15 min, washed and repeated 3 more times. Fmoc-glycine (3 eq, 513 mg, 1.65 mmol), HBTU (3 eq, 625 mg, 1.65 mmol), and DIEA (6 eq, 0.574 mL, 3.30 mmol) in DMF (15 mL) were added to the reaction flask and agitated for 2 h at ambient temperature and washed as before. The Fmoc group of glycine was deprotected, resin washed, and the Fmoc-glycine procedure was repeated twice more. After Fmoc deprotection, a solution of TFA/DCM (2:1, 20 mL) was added to the resin with agitation for 2 h at ambient temperature. The resin was filtered and resulting solution concentrated *in vacuo*. The residue was trituated with cold diethyl ether and purified using reverse phase HPLC using H_2_O/MeOH to yield **Fl-Tri(Gly_3_)**. The sample was analyzed for purity using a Shimadzu LC 2020 with a Phenomenex Luna 5µ C18(2) 100Å (30 x 2.00 mm) column; gradient elution with H_2_O/CH_3_CN.

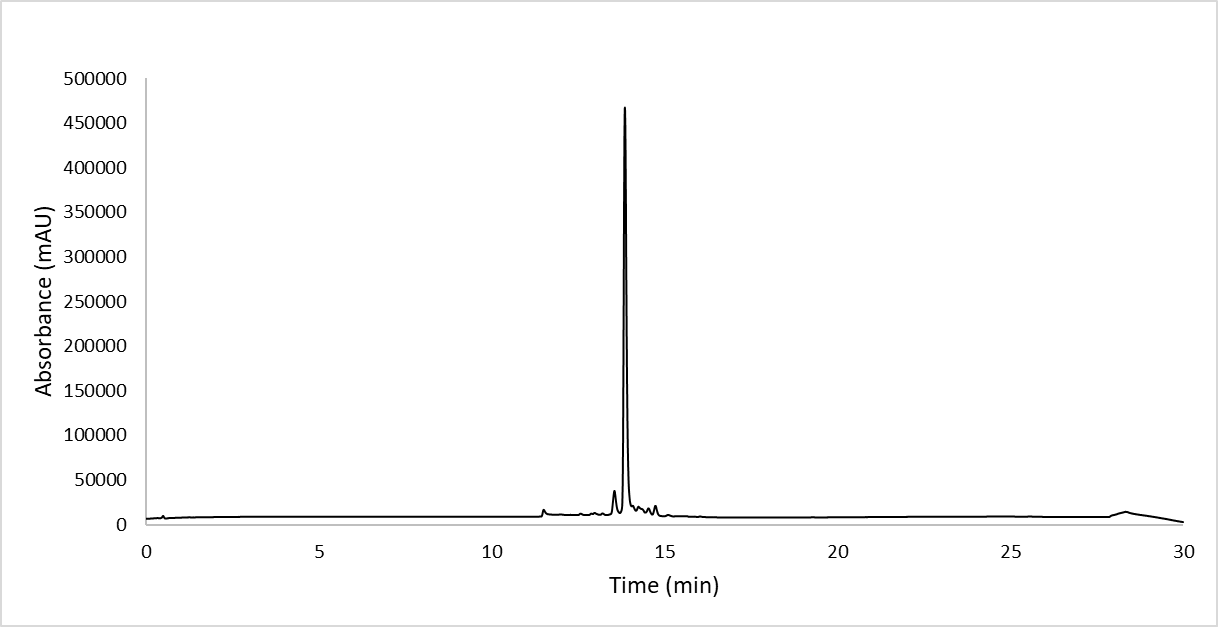

**Scheme S12. Synthesis of Fl-Tri(Gly_4_).**

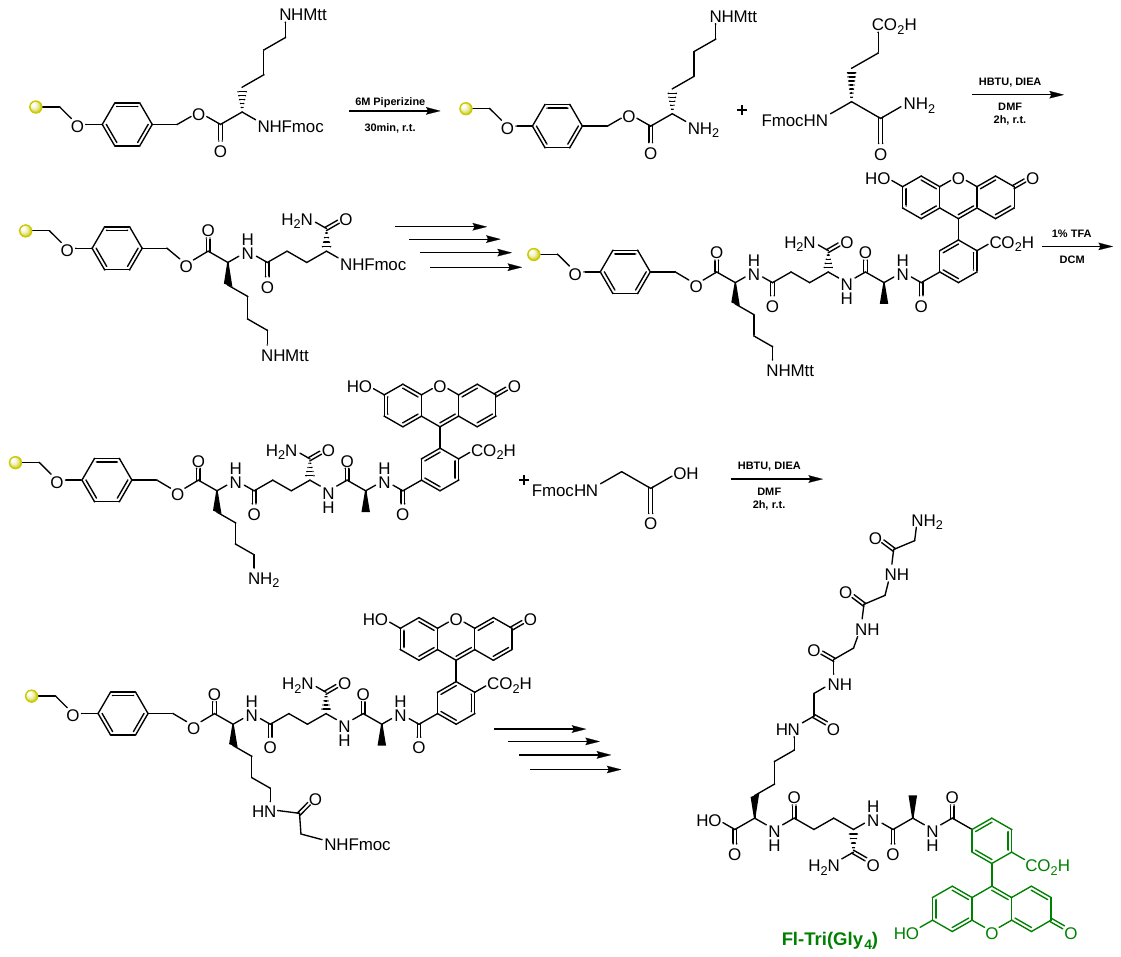

To a 25 mL peptide synthesis vessel charged with Fmoc-Lysine(Mtt)-Wang resin (1.0 g, 0.55mmol). The Fmoc protecting group was removed with 6 M piperazine/100 mM HOBt in DMF (15 ml) for 30 min at ambient temperature, then washed with washed with MeOH and DCM (3 x 15 mL each). Fmoc-D-glutamic acid α-amide (3 eq, 607 mg, 1.65 mmol), HBTU (3 eq, 625 mg, 1.65 mmol), and DIEA (6 eq, 0.574 mL, 3.30 mmol) in DMF (15 mL) were added to the reaction flask and agitated for 2 h at ambient temperature and washed as before. The Fmoc deprotection and coupling procedure was repeated as before using the same equivalencies with Fmoc-L-Alanine-OH. The Fmoc group of L-alanine was deprotected and resin coupled with 5(6)-carboxyfluorescein (2 eq, 413 mg, 1.1 mmol), HBTU (2 eq, 416 mg, 1.1 mmol) and DIEA (6 eq, 0.574 mL, 3.30 mmol) in DMF (15 mL) shaking overnight. The Mtt protecting group was removed by the addition of 1% TFA, 2.5% TIPS, in 10 mL DCM for 15 min, washed and repeated 3 more times. Fmoc-glycine (3 eq, 513 mg, 1.65 mmol), HBTU (3 eq, 625 mg, 1.65 mmol), and DIEA (6 eq, 0.574 mL, 3.30 mmol) in DMF (15 mL) were added to the reaction flask and agitated for 2 h at ambient temperature and washed as before. The Fmoc group of glycine was deprotected, resin washed, and the Fmoc-glycine procedure was repeated three more times. After Fmoc deprotection, a solution of TFA/DCM (2:1, 20 mL) was added to the resin with agitation for 2 h at ambient temperature. The resin was filtered and resulting solution concentrated *in vacuo*. The residue was trituated with cold diethyl ether and purified using reverse phase HPLC using H_2_O/MeOH to yield **Fl-Tri(Gly_4_)**. The sample was analyzed for purity using a Shimadzu LC 2020 with a Phenomenex Luna 5µ C18(2) 100Å (30 x 2.00 mm) column; gradient elution with H_2_O/CH_3_CN.

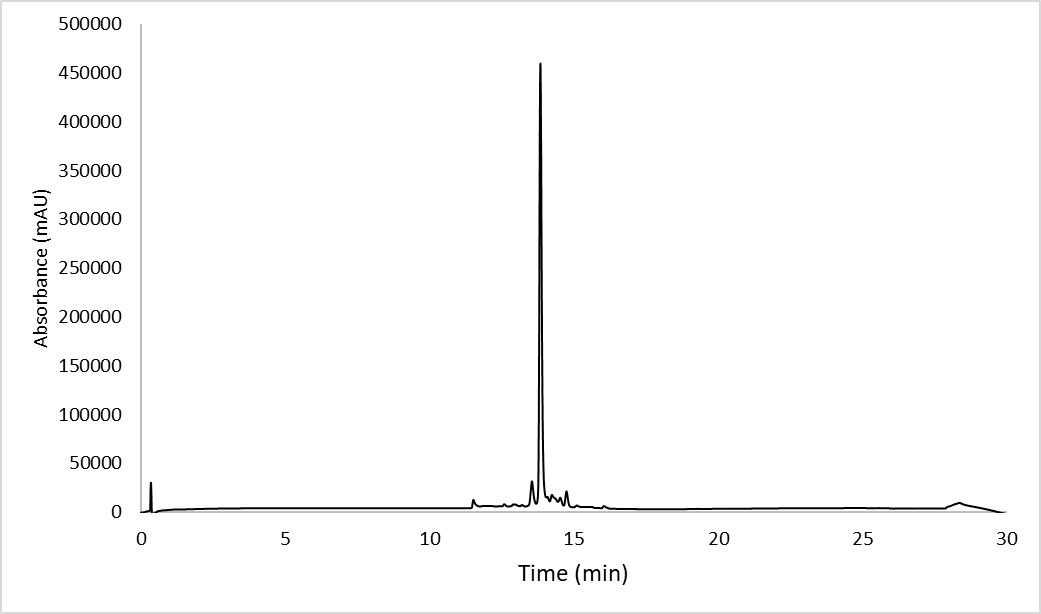

**Scheme S13. Synthesis of Fl-Tri(Gly_5_).**

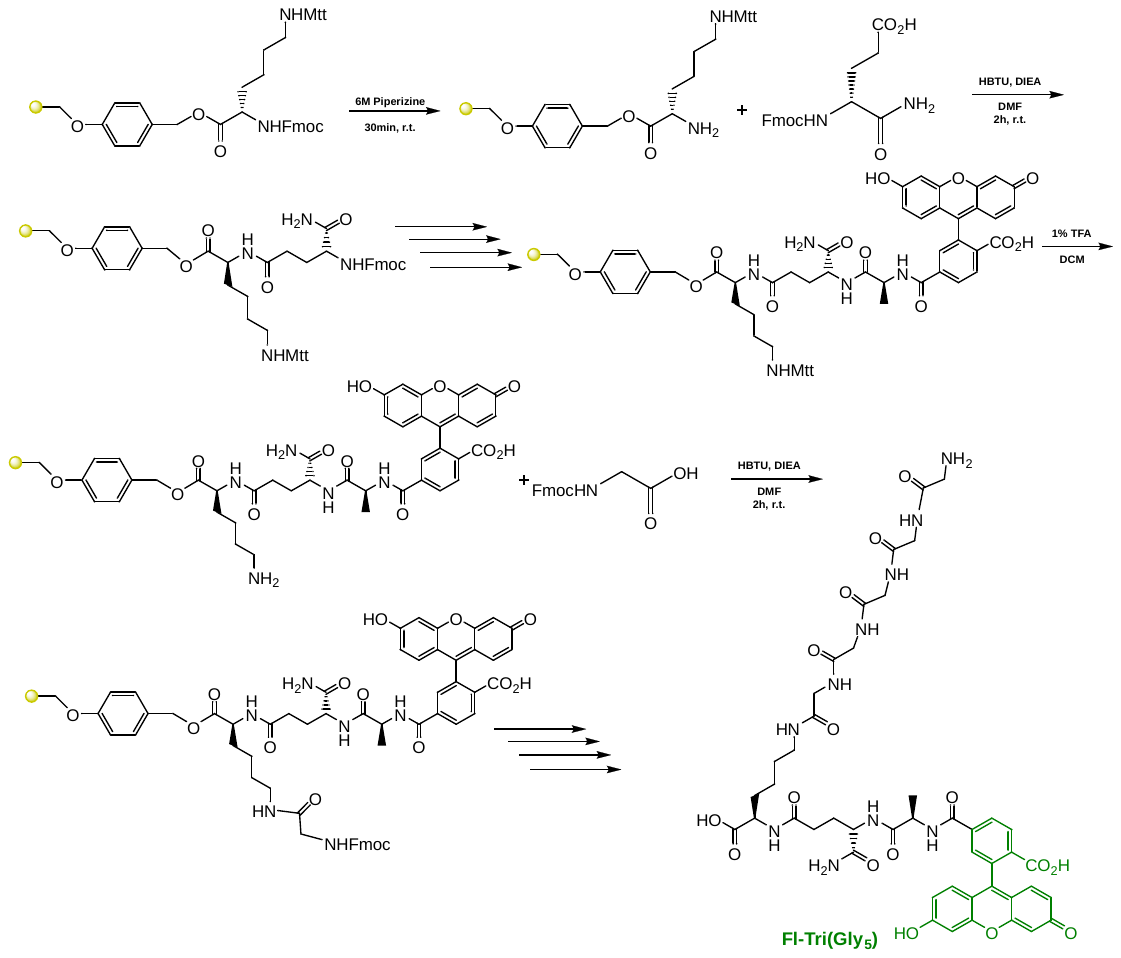

To a 25 mL peptide synthesis vessel charged with Fmoc-Lysine(Mtt)-Wang resin (1.0 g, 0.55mmol). The Fmoc protecting group was removed with 6 M piperazine/100 mM HOBt in DMF (15 ml) for 30 min at ambient temperature, then washed with washed with MeOH and DCM (3 x 15 mL each). Fmoc-D-glutamic acid α-amide (3 eq, 607 mg, 1.65 mmol), HBTU (3 eq, 625 mg, 1.65 mmol), and DIEA (6 eq, 0.574 mL, 3.30 mmol) in DMF (15 mL) were added to the reaction flask and agitated for 2 h at ambient temperature and washed as before. The Fmoc deprotection and coupling procedure was repeated as before using the same equivalencies with Fmoc-L-Alanine-OH. The Fmoc group of L-alanine was deprotected and resin coupled with 5(6)-carboxyfluorescein (2 eq, 413 mg, 1.1 mmol), HBTU (2 eq, 416 mg, 1.1 mmol) and DIEA (6 eq, 0.574 mL, 3.30 mmol) in DMF (15 mL) shaking overnight. The Mtt protecting group was removed by the addition of 1% TFA, 2.5% TIPS, in 10 mL DCM for 15 min, washed and repeated 3 more times. Fmoc-glycine (3 eq, 513 mg, 1.65 mmol), HBTU (3 eq, 625 mg, 1.65 mmol), and DIEA (6 eq, 0.574 mL, 3.30 mmol) in DMF (15 mL) were added to the reaction flask and agitated for 2 h at ambient temperature and washed as before. The Fmoc group of glycine was deprotected, resin washed, and the Fmoc-glycine procedure was repeated four more times. After Fmoc deprotection, a solution of TFA/DCM (2:1, 20 mL) was added to the resin with agitation for 2 h at ambient temperature. The resin was filtered and resulting solution concentrated *in vacuo*. The residue was trituated with cold diethyl ether and purified using reverse phase HPLC using H_2_O/MeOH to yield **Fl-Tri(Gly_5_)**. The sample was analyzed for purity using a Shimadzu LC 2020 with a Phenomenex Luna 5µ C18(2) 100Å (30 x 2.00 mm) column; gradient elution with H_2_O/CH_3_CN.

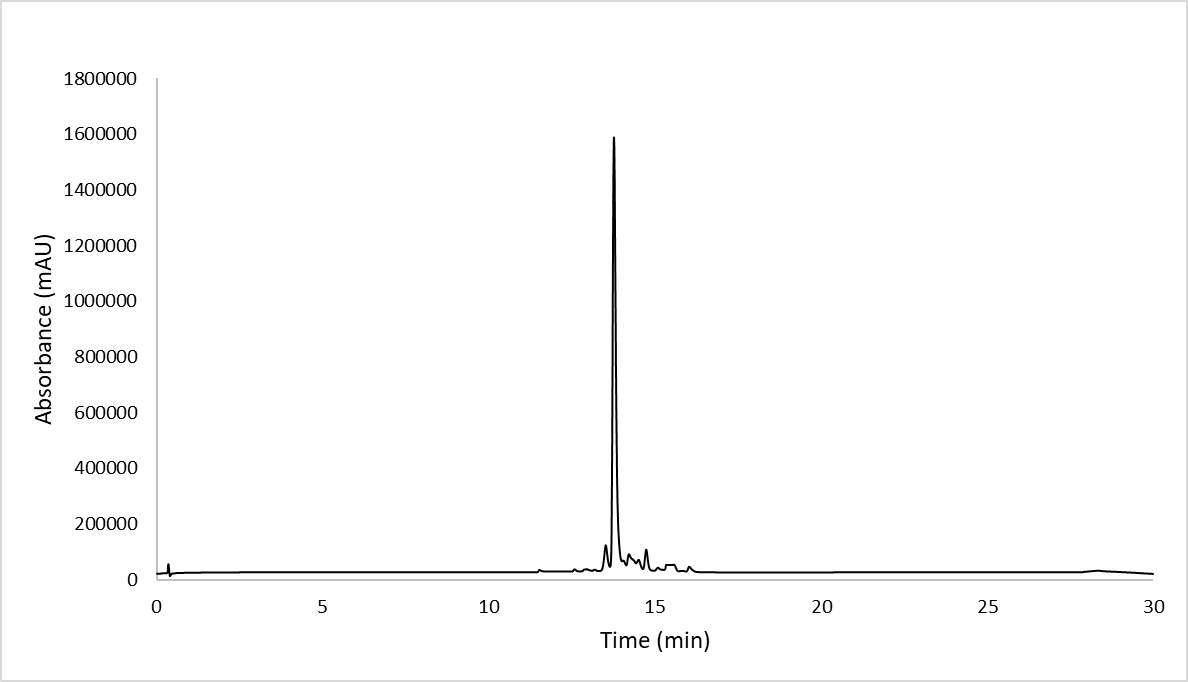

**Scheme S14. Synthesis of Fl-Tri(Gly_6_).**

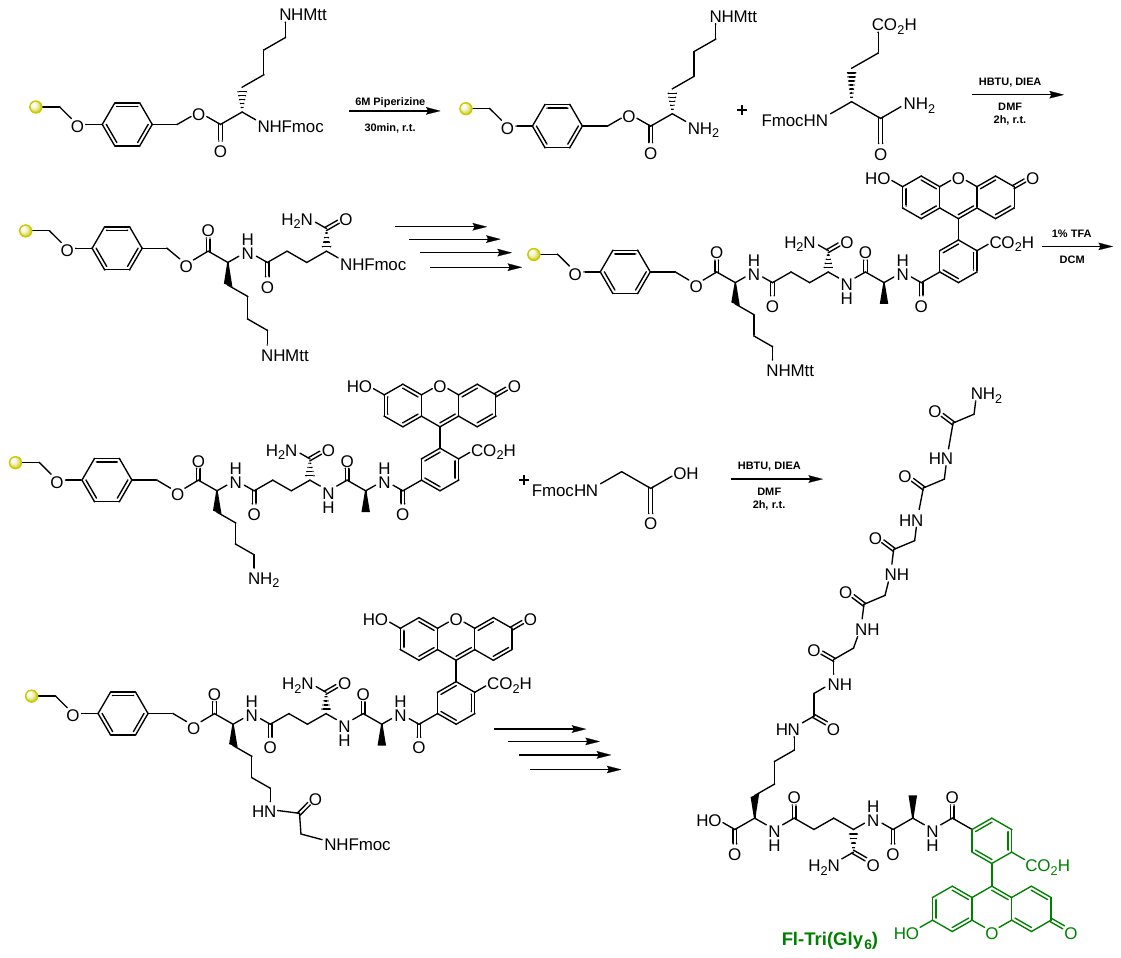

To a 25 mL peptide synthesis vessel charged with Fmoc-Lysine(Mtt)-Wang resin (1.0 g, 0.55mmol). The Fmoc protecting group was removed with 6 M piperazine/100 mM HOBt in DMF (15 ml) for 30 min at ambient temperature, then washed with washed with MeOH and DCM (3 x 15 mL each). Fmoc-D-glutamic acid α-amide (3 eq, 607 mg, 1.65 mmol), HBTU (3 eq, 625 mg, 1.65 mmol), and DIEA (6 eq, 0.574 mL, 3.30 mmol) in DMF (15 mL) were added to the reaction flask and agitated for 2 h at ambient temperature and washed as before. The Fmoc deprotection and coupling procedure was repeated as before using the same equivalencies with Fmoc-L-Alanine-OH. The Fmoc group of L-alanine was deprotected and resin coupled with 5(6)-carboxyfluorescein (2 eq, 413 mg, 1.1 mmol), HBTU (2 eq, 416 mg, 1.1 mmol) and DIEA (6 eq, 0.574 mL, 3.30 mmol) in DMF (15 mL) shaking overnight. The Mtt protecting group was removed by the addition of 1% TFA, 2.5% TIPS, in 10 mL DCM for 15 min, washed and repeated 3 more times. Fmoc-glycine (3 eq, 513 mg, 1.65 mmol), HBTU (3 eq, 625 mg, 1.65 mmol), and DIEA (6 eq, 0.574 mL, 3.30 mmol) in DMF (15 mL) were added to the reaction flask and agitated for 2 h at ambient temperature and washed as before. The Fmoc group of glycine was deprotected, resin washed, and the Fmoc-glycine procedure was repeated five more times. After Fmoc deprotection, a solution of TFA/DCM (2:1, 20 mL) was added to the resin with agitation for 2 h at ambient temperature. The resin was filtered and resulting solution concentrated *in vacuo*. The residue was trituated with cold diethyl ether and purified using reverse phase HPLC using H_2_O/MeOH to yield **Fl-Tri(Gly_6_)**. The sample was analyzed for purity using a Shimadzu LC 2020 with a Phenomenex Luna 5µ C18(2) 100Å (30 x 2.00 mm) column; gradient elution with H_2_O/CH_3_CN.

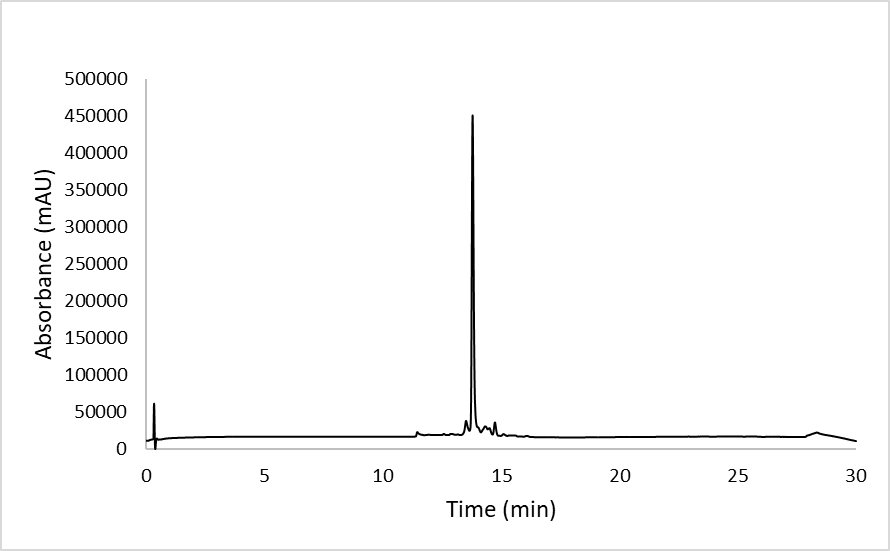

**Scheme S15. Synthesis of Coumarin-Tri(Asn).**

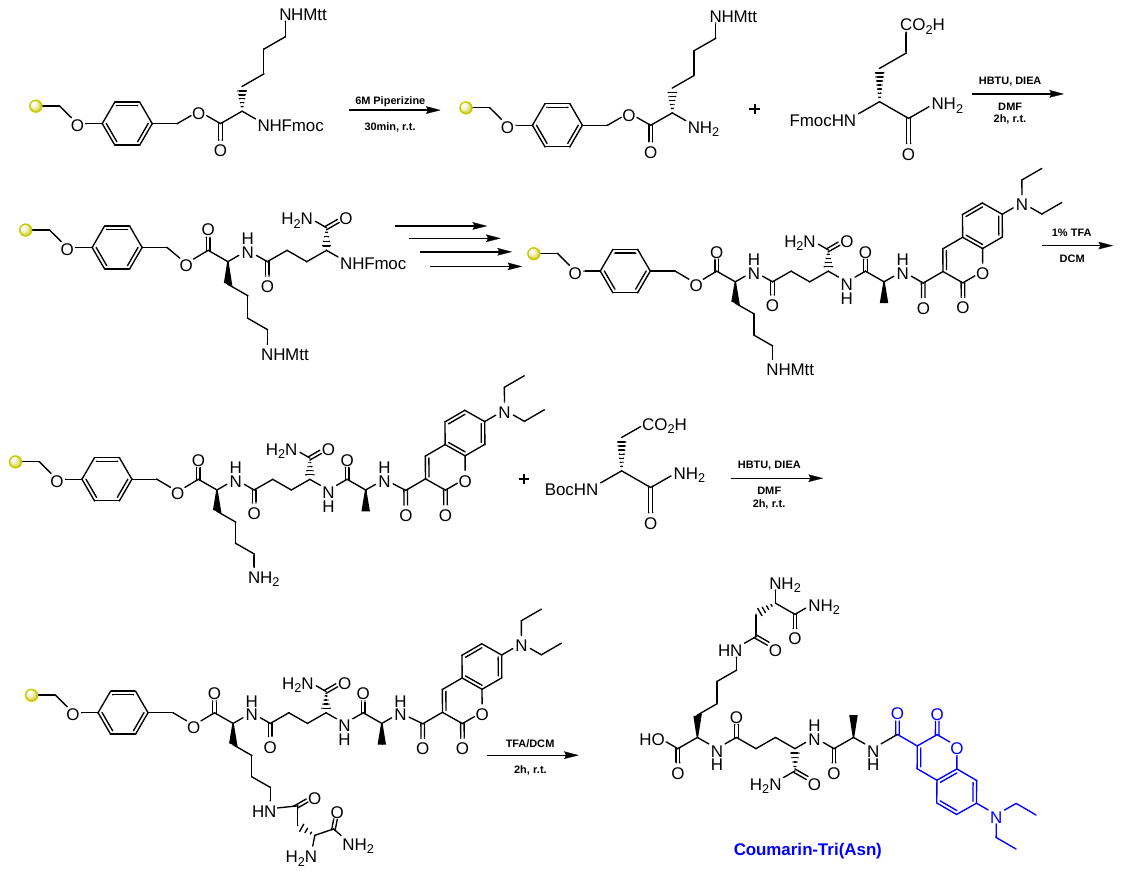

To a 25 mL peptide synthesis vessel charged with Fmoc-Lysine(Mtt)-Wang resin (1.0 g, 0.55mmol). The Fmoc protecting group was removed with 6 M piperazine/100 mM HOBt in DMF (15 ml) for 30 min at ambient temperature, then washed with washed with MeOH and DCM (3 x 15 mL each). Fmoc-D-glutamic acid α-amide (3 eq, 607 mg, 1.65 mmol), HBTU (3 eq, 625 mg, 1.65 mmol), and DIEA (6 eq, 0.574 mL, 3.30 mmol) in DMF (15 mL) were added to the reaction flask and agitated for 2 h at ambient temperature and washed as before. The Fmoc deprotection and coupling procedure was repeated as before using the same equivalencies with Fmoc-L-Alanine-OH. The Fmoc group of L-alanine was deprotected and resin coupled with 7-(Diethylamino)-2-oxo-2H-chromene-3-carboxylic acid (2 eq, 287 mg, 1.1 mmol), HBTU (2 eq, 416 mg, 1.1 mmol) and DIEA (6 eq, 0.574 mL, 3.30 mmol) in DMF (15 mL) shaking overnight. The Mtt protecting group was removed by the addition of 1% TFA, 2.5% TIPS, in 10 mL DCM for 15 min, washed and repeated 3 more times. Boc-D-aspartic acid α-amide (3 eq, 382 mg, 1.65 mmol), HBTU (3 eq, 625 mg, 1.65 mmol), and DIEA (6 eq, 0.574 mL, 3.30 mmol) in DMF (15 mL) were added to the reaction flask and agitated for 2 h at ambient temperature and washed as before. A solution of TFA/DCM (2:1, 20 mL) was added to the resin with agitation for 2 h at ambient temperature. The resin was filtered and resulting solution concentrated *in vacuo*. The residue was trituated with cold diethyl ether and purified using reverse phase HPLC using H_2_O/MeOH to yield **Coumarin-Tri(Asn)**. The sample was analyzed for purity using a Shimadzu LC 2020 with a Phenomenex Luna 5µ C18(2) 100Å (30 x 2.00 mm) column; gradient elution with H_2_O/CH_3_CN.

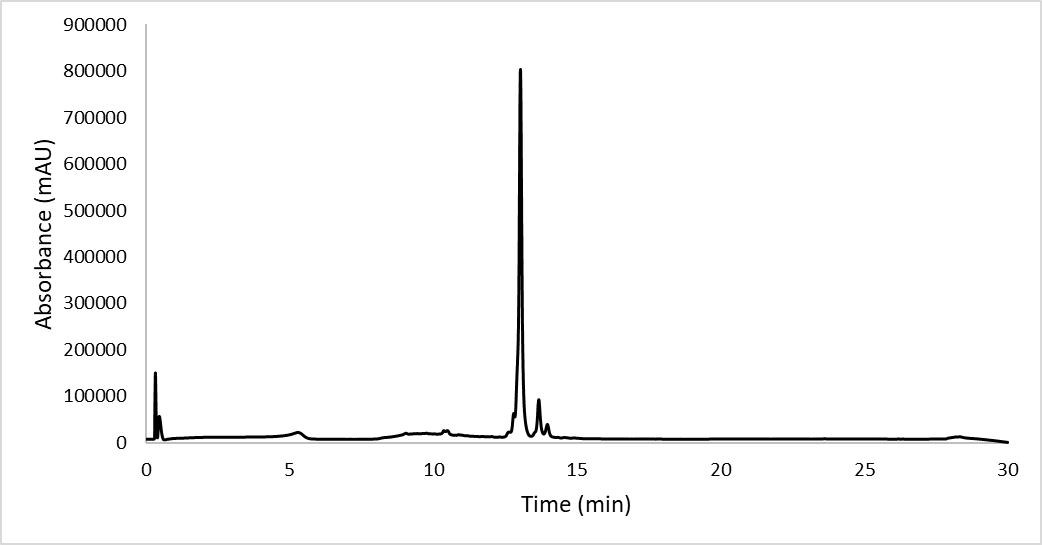

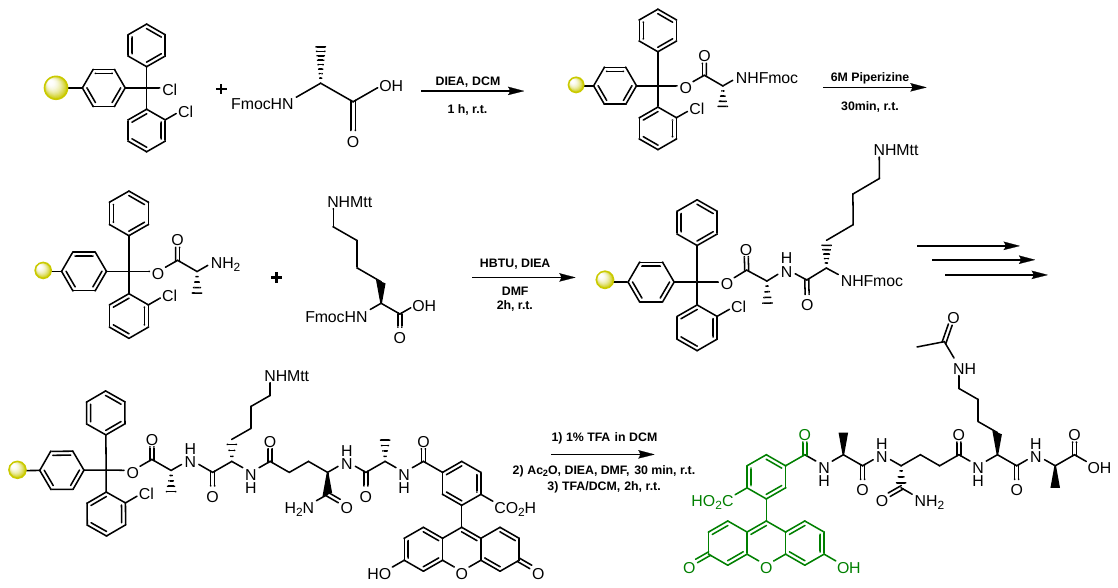

To a 25 mL peptide synthesis vessel charged with Fmoc-D-Alanine Wang resin (950 mg, 0.55 mmol). The Fmoc protecting group was removed with 6 M piperazine/100 mM HOBt in DMF (15 ml) for 30 min at ambient temperature, then washed with MeOH and DCM (3 x 15 mL each). Fmoc-L-Lys(Mtt)-OH (3 eq, 1.02 g, 1.65 mmol), HBTU (3 eq, 625 mg, 1.65 mmol), and DIEA (6 eq, 0.574 mL, 3.30 mmol) in DMF (15 mL) was added to the reaction flask and agitated for 2 h at ambient temperature. The Fmoc deprotection and coupling procedure was repeated as before using the same equivalencies with Fmoc-D-glutamic acid α-amide and Fmoc-L-alanine. The Fmoc group of L-alanine was deprotected and resin coupled with 5(6)-carboxyfluorescein (2 eq, 413 mg, 1.1 mmol), HBTU (2 eq, 416 mg, 1.1 mmol) and DIEA (6 eq, 0.574 mL, 3.30 mmol) in DMF (15 mL) shaking overnight. The resin was washed as before and added to a solution of 1% TFA / 5% TIPS in DCM and shaken for 10 min and washed. The step was repeated five times for removal of the Mtt group. Acetic anhydride (5 eq, 0.260 mL) and DIEA (10 eq, 0.956 mL) in DMF was added and resin shaken for 30 min at ambient temperature. The resin was washed and added to a solution of TFA/DCM (2:1, 20 mL) with agitation for 2 h at ambient temperature. The resin was filtered and resulting solution concentrated *in vacuo*. The residue was trituated with cold diethyl ether and purified using reverse phase HPLC using H_2_O/MeOH**.** The sample was analyzed for purity using a Shimadzu LC 2020 with a Phenomenex Luna 5µ C18(2) 100Å (30 x 2.00 mm) column; gradient elution with H_2_O/CH_3_CN.

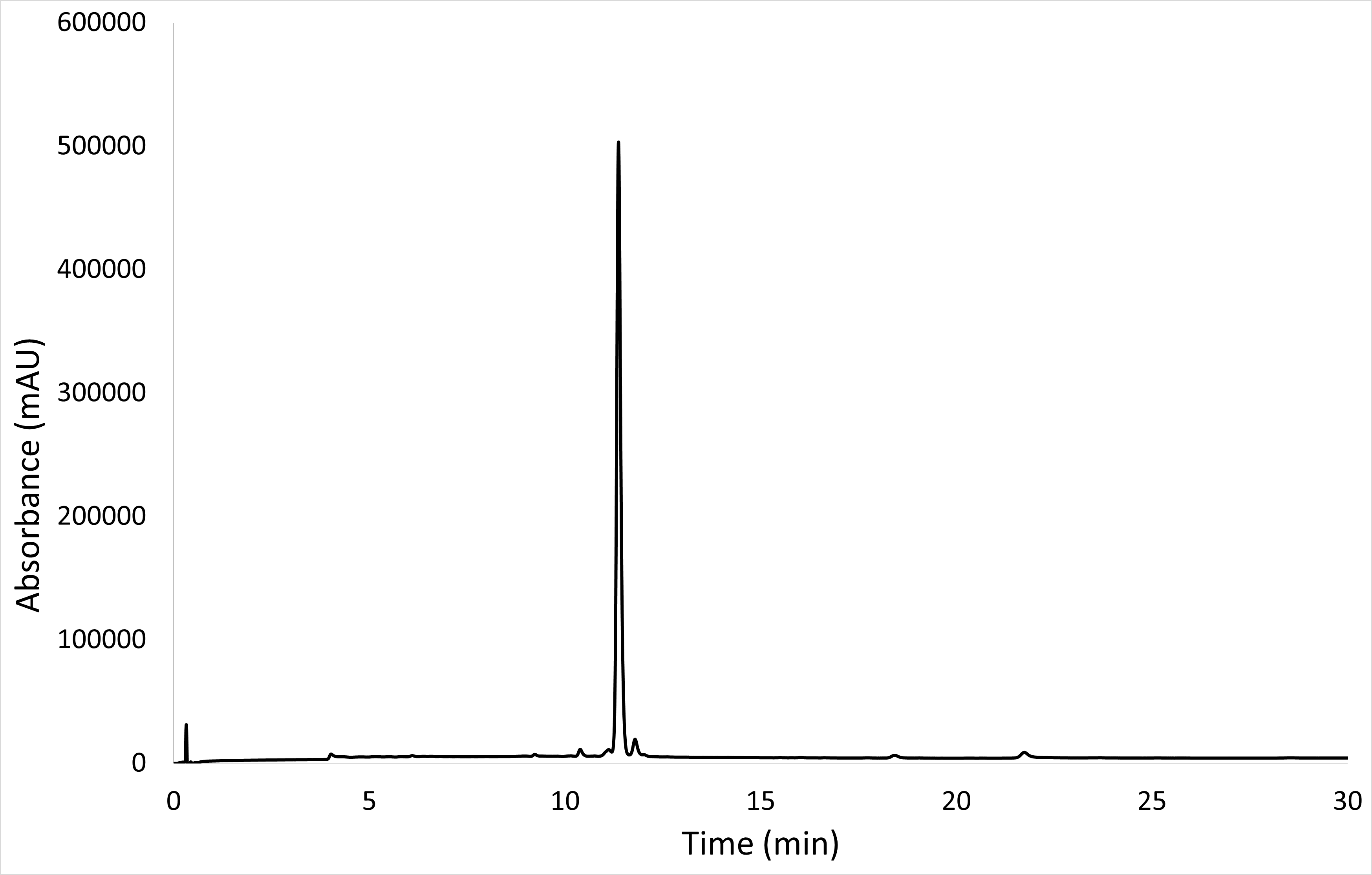

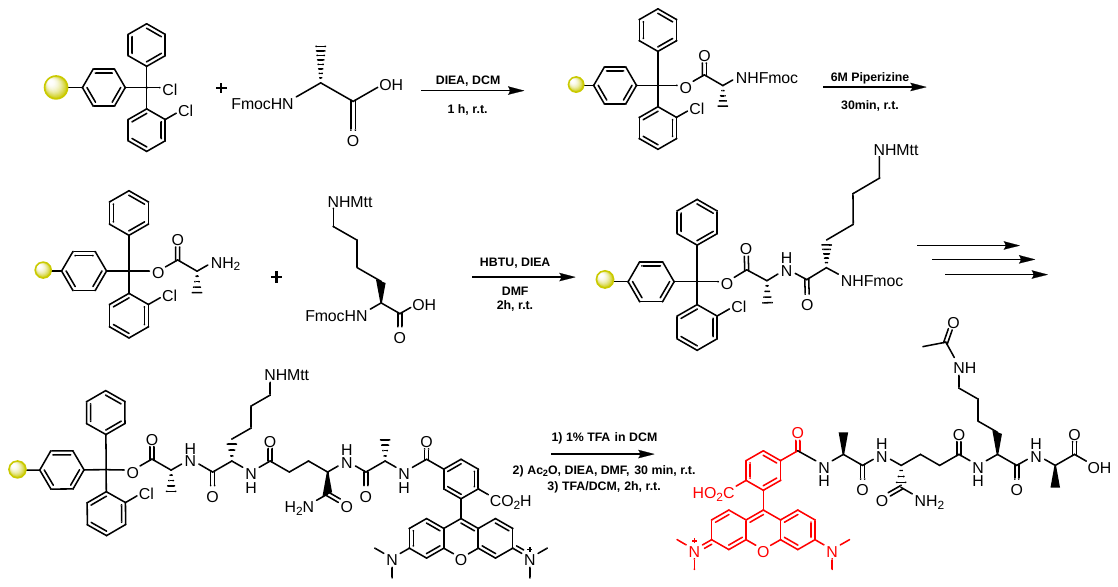

To a 25 mL peptide synthesis vessel charged with Fmoc-D-Alanine Wang resin (950 mg, 0.55 mmol). The Fmoc protecting group was removed with 6 M piperazine/100 mM HOBt in DMF (15 ml) for 30 min at ambient temperature, then washed with MeOH and DCM (3 x 15 mL each). Fmoc-L-Lys(Mtt)-OH (3 eq, 1.02 g, 1.65 mmol), HBTU (3 eq, 625 mg, 1.65 mmol), and DIEA (6 eq, 0.574 mL, 3.30 mmol) in DMF (15 mL) was added to the reaction flask and agitated for 2 h at ambient temperature. The Fmoc deprotection and coupling procedure was repeated as before using the same equivalencies with Fmoc-D-glutamic acid α-amide and Fmoc-L-alanine. The Fmoc group of L-alanine was deprotected and resin coupled with 5(6)-carboxytetramethylrhodamine (2 eq, 473 mg, 1.1 mmol), HBTU (2 eq, 416 mg, 1.1 mmol) and DIEA (6 eq, 0.574 mL, 3.30 mmol) in DMF (15 mL) shaking overnight. The resin was washed as before and added to a solution of 1% TFA / 5% TIPS in DCM and shaken for 10 min and washed. The step was repeated five times for removal of the Mtt group. Acetic anhydride (5 eq, 0.260 mL) and DIEA (10 eq, 0.956 mL) in DMF was added and resin shaken for 30 min at ambient temperature. The resin was washed and added to a solution of TFA/DCM (2:1, 20 mL) with agitation for 2 h at ambient temperature. The resin was filtered and resulting solution concentrated *in vacuo*. The residue was trituated with cold diethyl ether and purified using reverse phase HPLC using H_2_O/MeOH**.** The sample was analyzed for purity using a Shimadzu LC 2020 with a Phenomenex Luna 5µ C18(2) 100Å (30 x 2.00 mm) column; gradient elution with H_2_O/CH_3_CN.

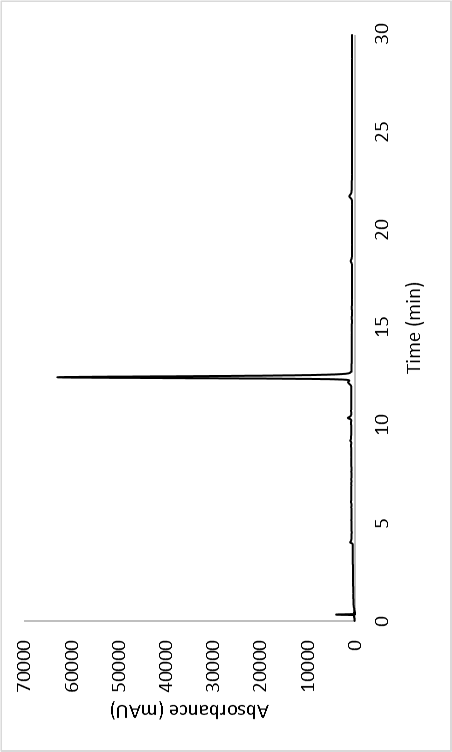

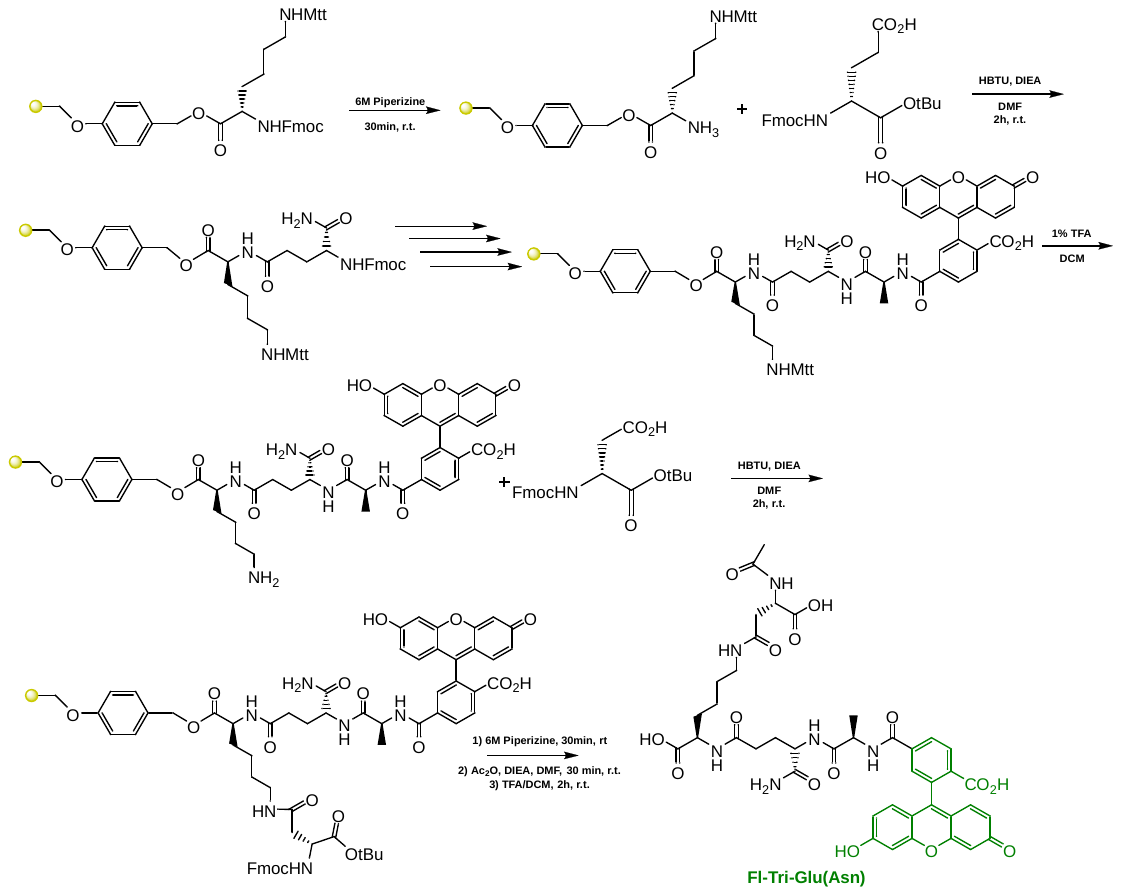
